# Pseudouridine-free Ribosome Exhibits Distinct Inter-subunit Movements

**DOI:** 10.1101/2021.06.02.446812

**Authors:** Yu Zhao, Jay Rai, Hongguo Yu, Hong Li

**Affiliations:** Institute of Molecular Biophysics, Florida State University, Tallahassee, FL 32306, USA; Biological Science Department, Florida State University, Tallahassee, FL 32306, USA; Department of Chemistry and Biochemistry, Florida State University, Tallahassee, FL 32306, USA

## Abstract

Pseudouridine, the most abundant form of RNA modification, is known to play important roles in ribosome function. Mutations in human DKC1, the pseudouridine synthase responsible for catalyzing the ribosome RNA modification, cause translation deficiencies and are associated with a complex cancer predisposition. The structural basis for how pseudouridine impacts ribosome function remains uncharacterized. Here we report electron cryomicroscopy structures of a fully modified and a pseudouridine-free ribosome from *Saccharomyces cerevisiae*. In the modified ribosome, the rearranged N1 atom of pseudouridine is observed to stabilize key functional motifs by establishing predominately water-mediated close contacts with the phosphate backbone. The pseudouridine-free ribosome, however, is devoid of such interactions and displays conformations reflective of abnormal inter-subunit movements. The erroneous motions of the pseudouridine-free ribosome may explain its observed deficiencies in translation.

## Introduction

Among all RNA that are known to be chemically modified, ribosomal RNA (rRNA) is only next to transfer RNA (tRNA) in modification abundance. Roughly 2% rRNA nucleotides contain 12 of the 140 known types of RNA modifications (Sharma and Lafontaine, 2015; Sloan et al., 2017). These modifications are believed to 1) impact rRNA stability (Gigova et al., 2014; Natchiar et al., 2017; Noon et al., 1998; Polikanov et al., 2015); 2) fine-tune ribosome function through modulating RNA-RNA and RNA-protein interactions (Natchiar et al., 2017; Polikanov et al., 2015); 3) prevent or promote ligand binding (Jack et al., 2011; Jiang et al., 2016; Natchiar et al., 2017); and 4) regulate ribosome assembly (Ghalei et al., 2017; Leppik et al., 2017; Liang et al., 2007, 2009b; Liu et al., 2008). Partial modification of rRNA nucleotides has recently been observed that can contribute to ribosome heterogeneity (Simsek et al., 2017). Both the diversity and the heterogeneity of modifications are believed to contribute to selective translation of specific mRNAs, adding a new layer of translational control of gene expression. Despite the rich biochemical and genetic evidence, the exact roles of the chemical modifications on rRNA on ribosome structure, however, have not been observed.

The most abundant modifications of rRNA are 2’-O-methylation and uridine isomerization (pseudouridylation), both of which are installed by small nucleolar ribonucleoparticles (snoRNPs) (Decatur and Fournier, 2003; Fatica and Tollervey, 2002; Maxwell and Fournier, 1995; Yu and Meier, 2014). The box C/D snoRNPs are responsible for site-specific addition of 2’-O-methyl groups at 80-100 sites in rRNA whereas the box H/ACA snoRNPs are responsible for isomerization of similar numbers of uridine. Each snoRNP is comprised of a catalytic subunit (fibrillarin as the methyltransferase and Cbf5p as the pseudouridine synthase in yeast, respectively), three accessory proteins, and a snoRNA capable of targeting the modification site(s) through base pairing (Decatur and Fournier, 2003; Liang and Li, 2011; Maxwell and Fournier, 1995; Watkins and Bohnsack, 2012). The effect of individual or a subset of snoRNP-mediated modification on cellular functions was previously studied by selectively removal of one or a small number of snoRNAs in yeast.(Babaian et al., 2020; Baxter-Roshek et al., 2007; King et al., 2003; Liang et al., 2007, 2009b; Piekna-Przybylska et al., 2008) Definitive defects in translation were observed in these yeast cells that can be attributed in part to changes in ribosome structures detected by dimethyl sulfate probing or the change in translation rate (Liang et al., 2007; Piekna-Przybylska et al., 2008).

In human, mutations or abnormal level of snoRNPs are associated with diseases ranging from reduced immunity against pathogens(Tiku et al., 2018) to predisposition to cancer (Heiss et al., 1998). Note that several special box C/D snoRNPs (U3, U14, U8) are essential components required for rRNA processing(Dragon et al., 2002) or as part of the telomerase ribonucleoprotein complex (Mitchell et al., 1999). which raises the question if a lack of chemical modification is the direct causes of these diseases. Understanding the snoRNPs-associated malignancy, therefore, requires dissecting the molecular mechanism of the molecules with which snoRNPs act upon or are associated under these conditions. Previously, mutation of the catalytic domain of Cbf5p, Asp95 in yeast, and *Dkc1^m^* in mammals, was shown to cause translation defects, most prominently in regulation of the IRES (internal ribosome entry site)-mediated, CAP-independent, translation without affecting the overall level of other genes (Bellodi et al., 2010; Jack et al., 2011; Yoon et al., 2006). This suggests a specific effect on ribosome in absence of pseudouridylation. Mutation of Cbf5 in human including *Dkc1^m^* is associated with Dyskeratosis Congenita (DC) that also has translational defects (Bellodi et al., 2010; Yoon et al., 2006). These findings underscore a direct impact of chemical modifications on ribosome-associated diseases.

Ribosome is a complicated molecular device visiting multiple states even when not translating and driven by thermal forces (Cornish et al., 2008; Frank and Gonzalez, 2010; Moore, 2012). Conformational dynamics lies at the heart of ribosome function optimally tuned for catalytic efficiency as well as translation fidelity. Translational factors and tRNA including their charge status promote conformational transitions required for the cyclic processes of initiation, elongation, and termination (Korostelev et al., 2008). Not surprisingly, factors that alter ribosome dynamics are also associated with translational deficiency (Sengupta et al., 2019; Sulima et al., 2014). Thus, characterization of these dynamic states provides a necessary biophysical basis for understanding ribosome function and associated diseases (Frank, 2016). While many of the conformational transitions, especially those of the bacterial ribosome, has been thoroughly characterized (Schmeing and Ramakrishnan, 2009). how chemical modifications influence these processes, and in turn, ribosome function, remain uncharacterized owing to the lack of structures of unmodified ribosome.

Here, we analyzed structures of vacant ribosomes isolated from yeast strains harboring the *nop1*-D243A or the *cbf5*-D95A mutation that is predicted to disrupt the chemical modification reactions catalyzed by fibrillarin and Cbf5p, respectively. We showed that while the *nop1*-D243A was not sufficient to remove 2’-O-methylation, the *cbf5*-D95A mutation resulted in ribosomes without pseudouridylation. Structure determination and comparison of the fully modified and the pseudouridine-free ribosome isolated from the *nop1*-D243A (2.57Å - 2.72Å) or the *cbf5*-D95A (2.89Å - 3.03Å) cells revealed a profound link between rRNA pseudouridylation and ribosome dynamics.

## Results

### Cell Growth and Polysome Profile Defects in nop1-D243A and cbf5-D95A Cells

Owing to the essentiality of fibrillarin and Cbf5p, we created yeast strains containing fibrillarin or Cbf5p mutants that lack the predicted catalytical residues. Fibrillarin belongs to the well-conserved 2’-O-methyltransferase family that include bacterial RrmJ(FtsJ) methyltransferase and the vaccinia virus VP39 methyltransferase whose catalytic residues had been identified by a combination of structural, bioinformatic and in vitro studies (Feder et al., 2003; Hager et al., 2002; Hodel et al., 1996) to contain a K-D-K-E tetrad. The tetrad corresponds to Lys150-Asp243-Lys272-Glu310 in yeast fibrillarin where Lys272 and Glu310 are believed to position the target RNA nucleotide while Lys150 and Asp243 play a catalytic role in the methyl transfer reaction. In addition, Glu198, similar to Asp83 of RrmJ, forms a close contact with the methyl donor, S-adenosyl-L-methionine (SAM), and is believed to be critical for SAM binding. Individual mutations of the tetrad residues, especially the aspartate residue, in RrmJ are detrimental to methylation both in vitro and in cells (Hager et al., 2002). Mutation of the aspartate in an archaeal fibrillarin also severely reduced in vitro methylation activity(Aittaleb et al., 2004). We, therefore, created the yeast strain whose genomic *nop1* encodes Asp243Ala (*nop1*-D243A). The mutation was confirmed by sequencing the PCR-amplified *nop1* locus from the genomic DNA of the strain (Supplementary Figure 1).

Similarly, the catalytic residues in Cbf5p were identified by its homology to the bacterial TruB pseudouridylase and by in vitro activity assays (Liang et al., 2009a). Critical to the isomerization reaction is a strictly conserved aspartate in all TruB family of pseudourylases (Liang et al., 2009a; Spedaliere et al., 2004) that corresponds to Asp95 of Cbf5p. We created a yeast strain whose genomic *cbf5* encodes Asp95Ala mutation (*cbf5*-D95A) and verified the strain by sequencing (Supplementary Figure 1). A similar strain was constructed previously with an episomal plasmid in *cbf5*-deletion cells, which resulted in viable cells that lack rRNA pseudouridylation (Jack et al., 2011; Zebarjadian et al., 1999).

Growth of the two mutant strains was assessed at three temperatures. In comparison with the wild-type, both strains exhibit severe growth deficiency at temperatures ranging from 25°C - 37°C (Figure 1). While the phenotype observed for *cbf5*-D95A is similar to that previously shown in the strain carrying D95A on an episomal plasmid, that observed for *nop1*-D243A is novel and suggests an important role of Asp243 of fibrillarin in cell growth.

**Figure 1.**
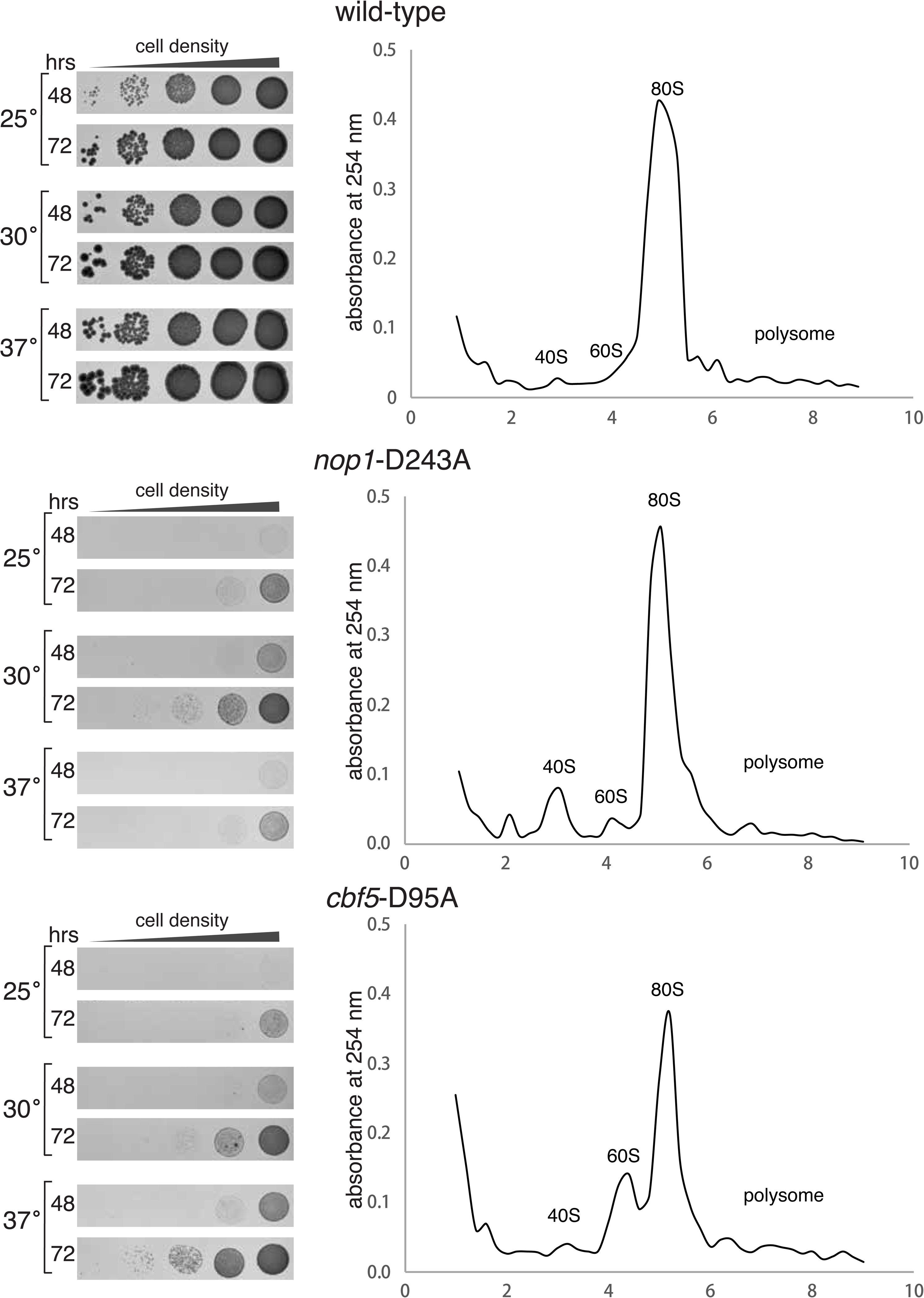
Growth and polysome profile comparison among the wild-type (top), *nop1*-D243A (middle), and *cbf5*-D95A (bottom) yeast cells. For growth analysis, cells were grown in a liquid YPD medium to saturation, washed twice and diluted with sterile water to OD of 1.0 (600 nm) before being spotted on solid YPD plate at each specified temperature. For polysome profile analysis, extracts from the cells grown to log phase were fractionated on a 10%-50% sucrose gradient and detected by UV absorbance at 254 nm.

To access the impact of the *nop1*-D243A and *cbf5*-D95A mutations in ribosome assembly, we performed sucrose density gradient centrifugation and compared polysome profiles between the wild-type and the two mutant cells grown in both normal and glucose depleted media. In comparison with the wild-type cells, there is an elevated level of free 40S in the *nop1*-D243A and that of free 60S in the *cbf5*-D95A cells, respectively, suggesting definitive but different defects in ribosome maturation in the mutant cells (Figure 1B). Despite so, both cells were able to produce sufficient 80S in supporting cellular functions (Figure 1B).

### A Fully Modified and a Pseudouridine-free Ribosome

We next examined the structures of the ribosome purified from the *nop1*-D243A and *cbf5*-D95A cells. We took advantage of the fact that slowly growing yeast accumulates inactive but homogenous ribosomes (Waldron et al., 1977) and purified them by two different methods to ensure consistent isolation of intact 80S from both strains (Materials and Methods) (Ben-Shem et al., 2011; Ben-Shem et al., 2010). We performed single particle analysis of the ribosomes isolated from the *nop1*-D243A and the *cbf5*-D95A cells, respectively. Initial classification and reconstruction revealed that a large majority of ribosome particles from either strain isolated by the affinity-based method are bound with the suppressor of target of Myb protein 1 (Stm1) (Figure 2), a known hibernation factor, that is similarly found in the inactive yeast 80S crystal structures (Ben-Shem et al., 2011; Ben-Shem et al., 2010). This form of the ribosome was used in comparative studies between the two ribosomes. 3D classification and focused refinement resulted in an overall resolution of 2.57-2.72 Å for the *nop1*-D243A ribosome (Supplementary Figure 2) and 2.89-3.03 Å for the *cbf5*-D95A ribosome (Supplementary Figure 3), respectively.

**Figure 2.**
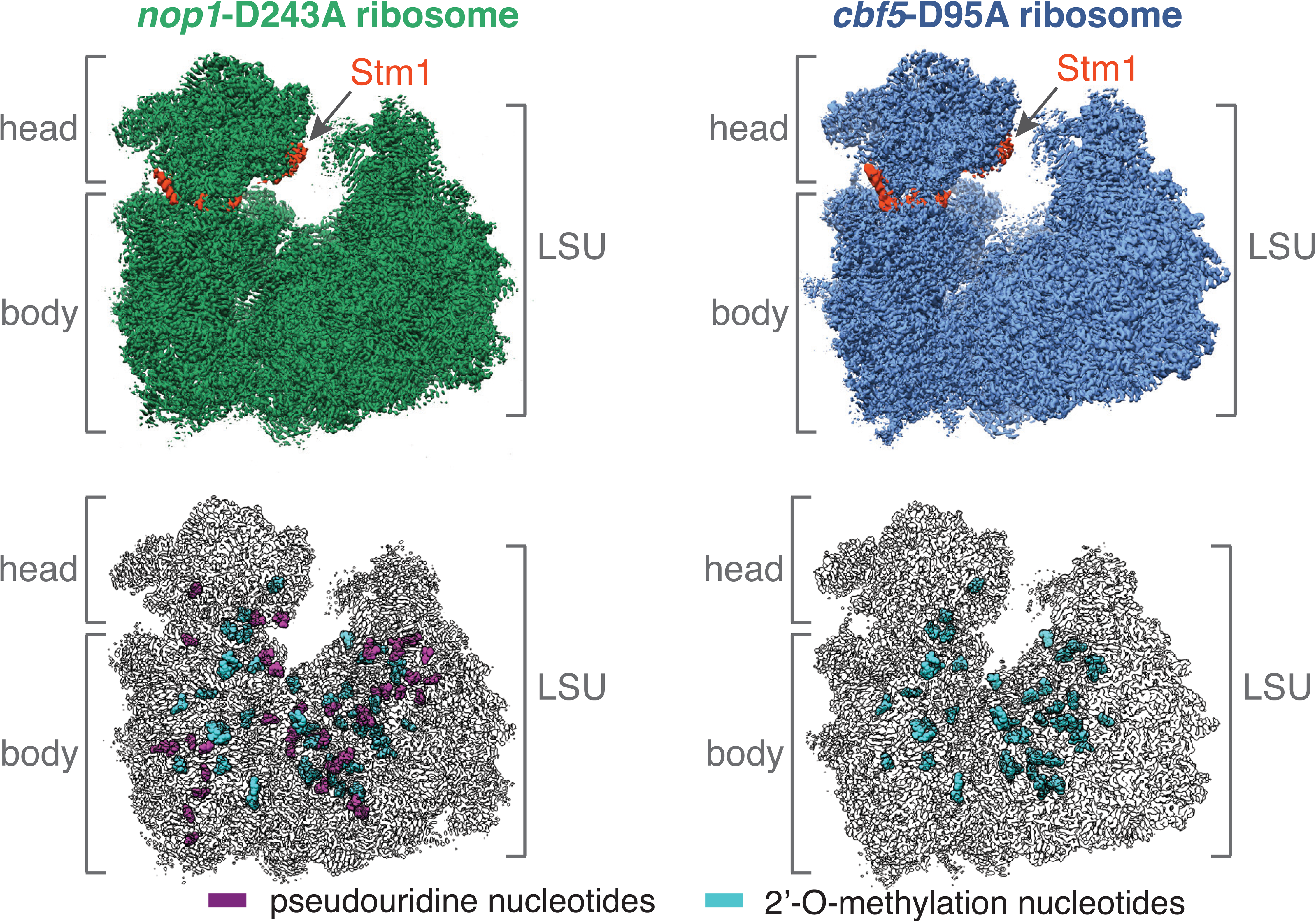
Overview of cryoEM structures of both the fully modified (*nop1*-D243A, green) and pseudouridine-free (*cbf5*-D95A, blue) yeast ribosome. “LSU” denotes the large subunit while “head” and “body” denote the head and body domains of the small subunit, respectively. The density belonging to the hibernating factor Stm1 is colored in red. Locations of pseudouridine and 2’-O-methylation are marked by purple and cyan, respectively.

The electron potential density readily revealed that, despite the known catalytic role of Asp243 of fibrillarin in 2’-O-methylation, the *nop1*-D243A ribosome still carries 2’-O-methylation at all positions targeted by snoRNPs (Supplementary Figure 4 & Table S1). Analysis of water molecules surrounding uridine showed that *nop1*-D243A ribosome display an interaction pattern consistent with the presence of pseudouridine at the predicted positions, as expected (Supplementary Figure 4). In contrast, the *cbf5*-D95A ribosome displays an interaction pattern indictive of a lack of pseudouridylation (Supplementary Figure 4 & Table S1), supporting the known catalytic role of Asp95 in isomerization. These structural observations suggest that *nop1*-D243A mutation is insufficient to remove 2’-O-methylation while *cbf5*-D95A mutation substantially reduced pseudouridylation. While the reason for the growth phenotype of *nop1*-D243A yeast strain is unclear, the high resolution of this ribosome provided us with an opportunity to model and analyze fully modified yeast ribosome (Figure 2). We were able to model 40 of the 45 pseudouridine, 54 of the 55 2’-O-methylation, and 11 of the 13 base modification nucleotides on the *nop1*-D143A ribosome (Supplementary Table 1). The structure of *cbf5*-D95A ribosome structure, on the other hand, allowed us to examine the structural features of the pseudouridine-free yeast ribosome (Figure 2) that was modeled with the same modified nucleotides except for pseudouridine (Supplementary Table 1).

### Distinct Motions of Ribosome with and without Pseudouridylation

We next analyzed and compared the structural conformations present in the fully modified and the pseudouridine-free ribosome that are bound with Stm1. For the best particles resulted from extensive selection, we performed 3D classification using a mask around the head of 40S followed by reconstruction. This procedure allowed us to examine conformationally distinct structures present in a single sample. Seven structures were obtained from the *nop1*-D243A ribosome with average resolutions of 2.8 Å to 3.2 Å (Figure 3A). Six structures resulted from the *cbf5*-D95A ribosome, albeit at lower resolutions of 4.3 Å to 5.8 Å (Figure 3B). To compare the conformations among the distinct ribosome structures, we computed and plotted the two rotation axes previously described as ribosome ratcheting and swiveling, respectively (Ling and Ermolenko, 2016). For a given pair of structures, we superimposed the densities of their large subunits onto the same 60S reference density and computed the angle and location of the transformation axis that brings one small subunit body to the other (ratcheting). We then aligned the small subunit bodies of both structures to the same reference body density and computed the angle and location of the transformation axis that brings one small subunit head to the other (swiveling). We also computed the same parameters for previously characterized ribosome conformations for comparison (Supplementary Figure 5).

**Figure 3.**
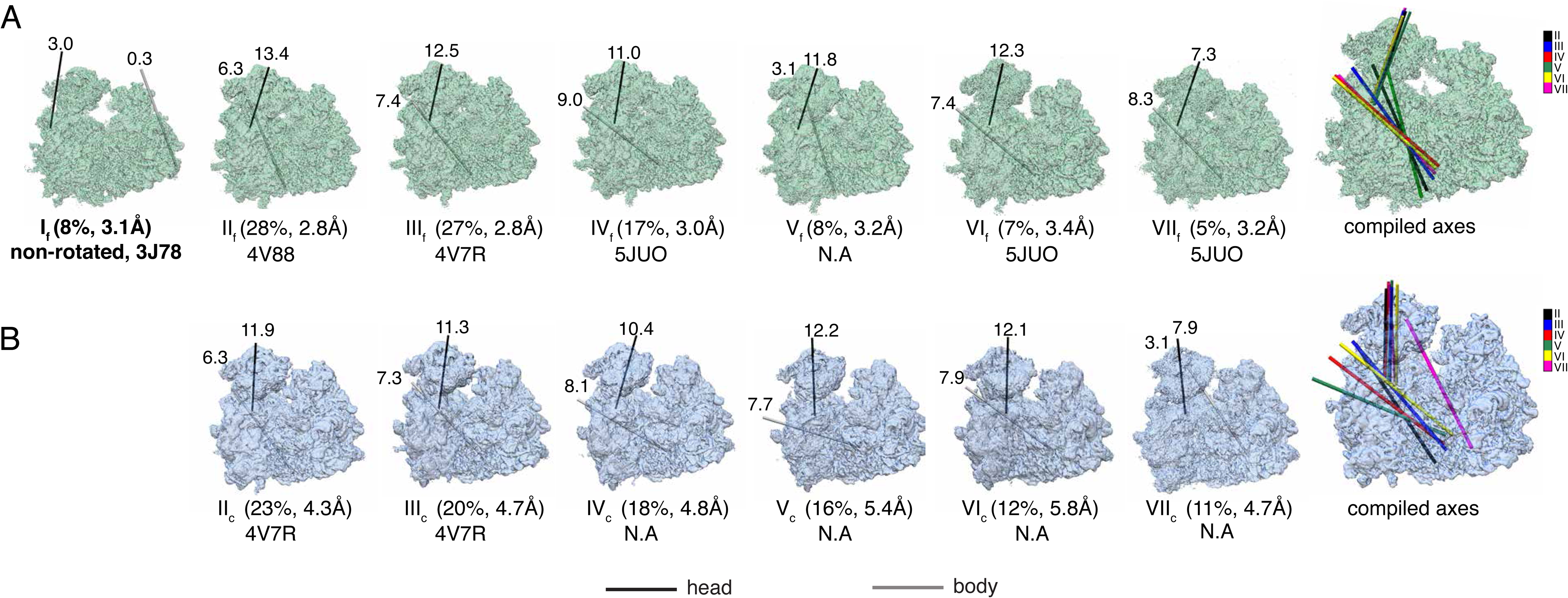
Comparison of ribosome conformations. Each classified and Stm1-bound ribosome conformation was compared with the non-rotated conformation (Structure I of *nop1*-D243A ribosome) except for Structure I of *nop1*-D243A that was compared with the previously known non-rotated structure (PDB ID: 3J78). The relationship between the two compared heads is indicated by the location and degree of rotation of the rotation axis in black (swivel). The relationship between the two compared bodies is indicated by the location and degree of rotation of the rotation axis in gray (ratchet). To reveal relative transformation relationships among the conformations, all rotation axes were also merged onto a single ribosome model and shown in different colors. Positive rotation is defined as clockwise when viewed into each rotation axis. The percentage of particles comprising each class over the total number of particles and the final resolution are shown in the parenthesis under each conformational class. **A.** Conformational distribution and relationships of the fully modified ribosome (*nop1*-D243A) structures. If a conformation is similar to a known ribosome structure, its PDB ID is indicated and if not, N.A. (not applicable) is indicated. **B.** Conformational distribution and relationships of the pseudouridine-free ribosome (*cbf5*-D95A) structures. If a conformation is similar to a known ribosome structure, its PDB ID is indicated and if not, N.A. (not applicable) is indicated.

Structure I of *nop1*-D243A ribosome (I_f_), which represents 8% of the total particles, closely matches the fully non-rotated conformation when bound with two tRNA and a model mRNA molecules (PDB ID: 3J78) (Svidritskiy et al., 2014) (Figure 3A). Therefore, we compared the six remaining structures with respect to I_f_. The remaining structures resemble a series of pre-translocation intermediates with ratcheting angles ranging 3°-9° and swiveling angles ranging 7°-13°. None of these, however, resembles the conformation of the fully rotated conformation (PDB ID: 3J77) captured when bound with one tRNA at its P/E site with ∼11° ratcheting and near 0° swiveling (Svidritskiy et al., 2014). Among the intermediates, structures II_f_ and III_f_ that together represent 55% of the total particles, resemble the two conformations captured in the crystal structures (PDB IDs: 4V88 and 4V7R) (Ben-Shem et al., 2011; Ben-Shem et al., 2010) based on the body rotation. These two conformations differ in ratcheting by ∼5° and swiveling by ∼7° (Supplementary Figure 5). Interestingly, structures IV_f_, VI_f_, and VII_f_ that together represent 29% of the particles, are highly similar among themselves and resemble the pre-translocation intermediate observed when the Taura Syndrome Virus IRES, eEF2-GDP are bound to ribosome with a fully rotated 40S body (PDB ID: 5JUO) (Abeyrathne et al., 2016) (Supplementary Figure 5). Structure V_f_, that comprises 8% of the particles has a nearly non-rotated body but swiveled head of 40S. Strikingly, all rotations of the *nop1*-D243A ribosome are anchored on two sites within the 40S subunit, one for ratcheting and one for swiveling, respectively, suggesting that *nop1*-D243A ribosome is shaped to support these intrinsic motions (Figure 3).

3D classification of the *cbf5*-D95A ribosome resulted in six structures. Unlike *nop1*-D243A ribosome, the *cbf5*-D95A ribosome do not include the non-rotated conformation. Like *nop1*-D243A ribosome, no *cbf5*-D95A ribosome conformation matches the fully rotated conformation (3J77). Therefore, the *cbf5*-D95A ribosome was compared to the non-rotated structure I_f_ of *nop1*-D243A ribosome. With exception to structures II_c_ and III_c_ that together make up 43% of the particles and somewhat resemble the crystal structure 4V7R, none matches known ribosome conformations we examined (Supplementary Figure 5). Furthermore, these rotations, especially ratcheting, do not anchor on single sites within ribosome unlike those of the *nop1*-D243A ribosome (Figure 3B).

To examine the major modes of motion among all particles, we performed multi-body analysis on ribosome from each cell by dividing ribosome into three rigid bodies: the head, the body, and the large subunit. The resulting top three normal mode motions from each ribosome were compared in terms of the direction and the range of the motions. While the major motion of *nop1*-D243A ribosome is that the head and body of 40S rotate in sync with respect to 60S (Supplementary Figure S6 & Supplementary data movie 1), that of *cbf5*-D95A ribosome comprises of independent rotations of the head and body of 40S with respect to 60S in different directions (Supplementary Figure S6 & Supplementary data movie 2). The next two major eigen vectors also differ in the degree and direction of motions substantially from those of the *nop1*-D243A ribosome (Supplementary Figure S6 & Supplementary data movies 3-6). This result further supports that the conformational dynamics differ largely between the fully modified and the pseudouridine-free ribosome.

### Pseudouridine Clusters Stabilize Key Motion Elements

Mapping the positions of modified nucleotides onto the refined yeast 60S structure model readily revealed that modifications are clustered within three functional centers: the peptidyl transferase center (PTC), the inter-subunit bridge (ISB) and the A-site finger (ASF) (King et al., 2003). Notably, biochemical analyses showed that mutations in proteins closely associated with PTC, rpL10p and rpL3p (Meskauskas and Dinman, 2008; Sulima et al., 2014), had profound effects on ribosome dynamics and thus translation fidelity. To illustrate how these modification clusters are linked to the mechanics of ribosome motion, we computed pair-wise displacements of the 25S rRNA nucleotides between the non-rotated and the rotated structures when their 40S subunits are superimposed (Figure 4A) and mapped the sites of psedouridylation and 2’-O-methylation on the nucleotide-wise displacement plot. Strikingly, pseudouridine clusters associated with PTC and ISB largely coincide with their minimal motion while the pseudouridine cluster associated with ASF experiences a significant translocation. This result suggests a motion-dependent distribution of pseudouridine on 60S.

**Figure 4.**
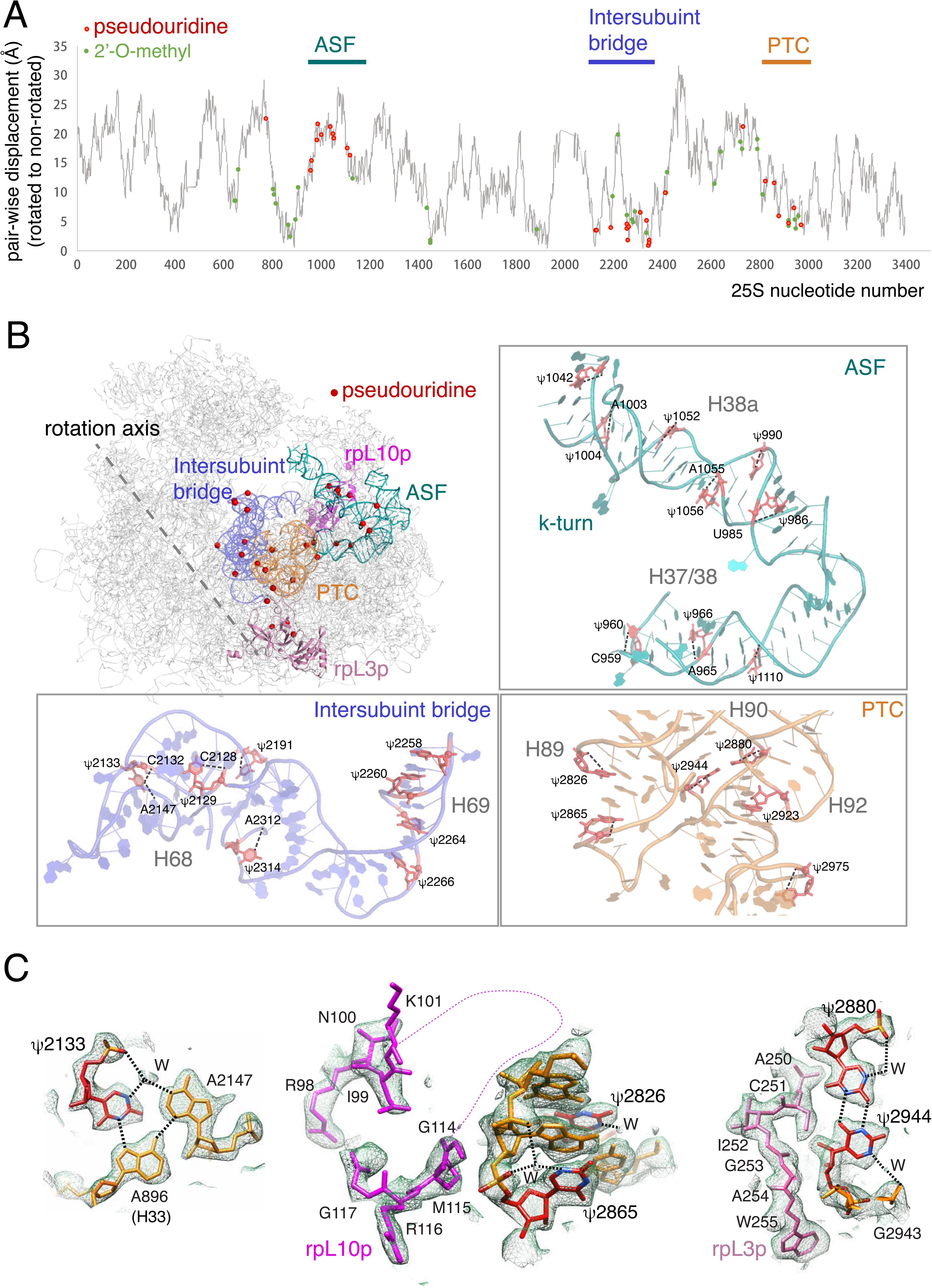
Observed modifications in key regions important to ribosome motions. **A.** Pair-wise displacement of the 25S rRNA from the rotated (PDB ID: 3J77) to the non-rotated transformation (PDB ID: 3J78) superimposed with sites of 2’-O-methylation (green) and pseoduridylation (red). The modification-rich, especially pseudouridine-rich, clusters coincide with three functional regions, the A-site figure (ASF, dark green bar), the intersubunit bridge (ISB, blue bar), and the peptidyl transfer center (PTC, orange bar). **B.** Observed pseudouridine distributions in ASF, ISB, and PTC, of the *nop1*-D243A ribosome and in relation to the rpL10 (magenta) and rpL3 (pink) proteins. The rotation axis marks the major location of ratchet motion in fully modified ribosome. Insets illustrate the observed intra-strand interactions by pseudouridine within the three clusters. Dashed lines depict water-mediated contacts of the N1 atom of a pseudouridine to the phosphate sugar backbone of either self or adjacent nucleotides. Pseudouridine nucleotids not associated with dashed lines indicate weak or absence of density supporting the interactions. **C.** Examples of how pseudouridine in ASF (ψ2133) and PTC (ψ2826 and ψ2865, ψ2880, and ψ2944) participate in networks of hydrogen bonds observed in the *nop1*-D243A ribosome. Dash lines represent the residues 101-113 not modeled for rpL10p due to lack of density.

Distribution of pseudouridine on the 18S is more scattered and does not seem to be correlated with the rotated-to-non-rotated displacement of nucleotides (Supplementary Figure 7). On the other hand, when modification sites are mapped on to the nucleotide-wise displacement plot from the swivel to the non-swivel 18S structures, it is clear that helix 44 (H44) is stationary, although it completely lacks chemical modifications. On the other hand, H44 interacts closely with the conserved helix 69 (H69) of ISB that contains a pseudouridine cluster (Supplementary Figure 7), suggesting that modifications on H69 may also impact head movement.

Examination of interactions mediated by pseudouridine in the *nop1*-D243A ribosome structure indicates that pseudouridine primarily stabilizes intra-strand interactions while also participate in other interactions. The density peak near the N1 atoms of a large number of pseudouridine that likely represents a water molecule is observed to be within hydrogen bond distance to the non-bridging oxygen of phosphate backbone of either the same or the preceding nucleotide (Figure 4 & Supplementary Figure 4). The same features are absent in the *cbf5*-D95A ribosome (Supplementary Figure 8). The intra-strand stabilization helps to maintain the integrity of the pseudouridine-rich helix 38a in ASF during translocation (Figure 4). Within ISB, we observed the same type of intra-strand interactions such as those in helix 68 (H68) as well as base-triple interactions such as those among Ψ2133, A2147 of H68 and A896 of helix 33 (H33) (Figure 4C). The pseudouridine-stabilized ISB interacts with the stationary H44 of the 40S, which could facilitate head swivel (Supplementary Figure 7).

Interestingly, pseudouridine-mediated interactions within PTC coincide with the large subunit proteins previously shown to impact ribosome dynamics (Meskauskas and Dinman, 2008; Sulima et al., 2014). The P site loop of rpL10p, 102-112, resides near a pair of pseudouridine, Ψ2826 and Ψ2865 of helix 89 (H89), that participate in intra-strand hydrogen bonds (Figure 4C). The highly charged N-terminus and a long loop (236-260) of rpL3p sandwich another pair of PTC pseudouridine, Ψ2880 and Ψ2944 of helix 90 (H90), that forms cross-strand interactions uniquely stabilized by pseudouridine (Figure 4C). These protein elements may further strengthen or counterbalance those established by pseudouridine at PTC as ribosome undergoes the non-rotated to the rotated transitions.

## Discussions

To examine the functional roles of rRNA modifications, we constructed yeast strains harboring either the *nop1*-D243A or the *cbf5*-D95A genomic mutation that is predicted to inactivate the respective modification enzymes while maintaining proper snoRNP assembly. We showed by structural biology methods that while the 80S ribosome from the *nop1*-D243A yeast contains fully modified rRNA, which indicates an active fibrillarin D243A, that from the *cbf5*-D95A yeast lacks pseudouridine. We compared structures, both global and local, as well as the dynamics between the *nop1*-D243A and the *cbf5*-D95A ribosome. While local structural environments around modified sites are similar at the current structure resolutions, the overall structural dynamics of the three major domains of 80S exhibits large differences between the fully modified and the pseudouridine-free 80S.

A large body of literature has eluted to a critical role of rRNA pseudouridylation in ribosome maturation and translation. Selective removal of individual or a subset of pseudouridine (Babaian et al., 2020; King et al., 2003; Liang et al., 2007; Piekna-Przybylska et al., 2008) caused an accumulatively negative effect on cell growth. These cells showed defective rRNA processing and altered ribosome profiles. The use of dual plasmid luciferase assay permitted direct assessment of translation in yeast cells (Ghalei et al., 2017; Rakauskaite et al., 2011). In the *cbf5*-D95A cell where rRNA pseudouridylation was abolished, decreased fidelity and defects in initiations were observed (Jack et al., 2011), suggesting a change of ribosome structural properties. Interestingly, DKC1 mutations in both mouse cells and the Dyskeratosis Congenita patient cells differentially impact the CAP-dependent and the IRES-mediated translation.(Bellodi et al., 2010; Yoon et al., 2006) Our structural analysis of the fully modified and the pseudouridine-free ribosome provides a structure-based explanation for the observed translation defects in cells. We believe that the three pseudouridine clusters within the large subunit stabilize the key regions that are necessary for proper inter-subunit movements. In absence of pseudouridine, ribosome losses the ability to drive the motions accurately.

Our analysis of the pseudouridine-free ribosome dynamics does not eliminate other possible deleterious effects in these ribosome such as changes in ligand binding. These effects, when combined with altered ribosome movements, whether directly or indirectly, can have differential impacts on different translation processes. Ψ2919, for instance, base pairs with the 3’ end of A site tRNA and the loss of its modifying snoRNA, snR10, led to reduced rate of translation (King et al., 2003). Similarly, loss of snoRNAs that guide pseudouridylation of those on H38 of ASF exhibited increased sensitivity to several antibiotics binding even though these antibiotics do not directly bind to ASF (Piekna-Przybylska et al., 2008). Finally, the hyper modified m(1)acp(3)Ψ1191 (U1248 in human) contributes to P site decoding interactions and its loss reduced the level as well as the activity of ribosome in yeast(Babaian et al., 2020; Liang et al., 2009b). In human, loss of the hypo-modification of U1248 is a prominent feature in multiple forms of cancer(Babaian et al., 2020).

The Stm1-bound ribosome represents an inactive form of the ribosome that accumulates under cell stress or carbon depletion (Ashe et al., 2000). The observed structural properties of the Stm1-bound ribosome do not reflect all conformations ribosome visits throughout its functional cycle. However, it is a landing-pad for yeast ribosome structural studies owing to its sample homogeneity, and has been discovered to mimic the rotated, mRNA-bound form of the 80S. Thus, the modes of ribosome motion can be representative of those possible in actively translating ribosome. Though this study focuses on structural comparison of a single ribosome form, similar differences are likely to be observed between the fully modified and the pseudouridine-free ribosome in other states, if these ribosome states are to be assembled and analyzed structurally in the future.

## Experimental Methods

### Construction of Yeast Strains

*Saccharomyces cerevisiae* haploid strain BY4741 was used as the genetic background to create the mutant strains by applying the Polymerase Chain Reaction (PCR) based method (Gardner and Jaspersen, 2014). In brief, a gene-specific mutation module was PCR amplified using pFA6a-3HA-kanMX6a plasmid as template and was later transformed into the wild-type cells. Transformants were selected with geneticin (G418). Correct mutation was confirmed by colony PCR and DNA sequencing. For both *nop1*-D243A and *cbf5*-D95A, the aspartic acid was mutated to alanine by changing the codon from GAT to GCT.

Growth phenotype of mutant strains was characterized by serial dilution and spotting assay analysis. For polysome profile analysis, cells were grown to log phase (OD600 0.4-0.6) in 200ml of liquid yeast peptone dextrose (YPD) before being shifted to YPD or yeast peptone (YP) for 10 minuets. Cells were harvested at 5000 rpm for 20 minutes in presence of 100 μg/ml of cycloheximide (CHX) followed by washing and resuspending with profiling buffer (10mM Tris-HCl pH7.5, 100mM KCl, 5mM MgCl_2_, 6mM β-mercaptoethanol (β-ME), 100 μg/ml CHX). Cells were broken by glass bead beating for six sets of 20sec-vertexing with 40sec-intervals at 4 °C. Lysates were cleared by centrifugation at 16000rpm in a tabletop centrifuge for 20 minutes at 4 °C. 9-11 UV 260nm units of the cleared lysate were loaded onto a 10%-50% sucrose gradient and centrifuged for 2.5 hours in a sw41Ti rotor at 4 °C. Fifty 200μl-fractions were manually taken from top to bottom of the gradient. UV absorbance at 254 nm was measured for each fraction and plotted against volume to generate the polysome profiles (Figure 1).

### Purification of Hibernating Ribosomes

Inactive ribosomes were purified using the standard sucrose density gradient ultracentrifugation method. We used the same *S. cervisiae* BY4741 strains harboring either *nop1*-D243A or *cbf5*-D95A that also contain a streptavidin-binding peptide tag on Kre33p, TAP tag on Pwp2p. Yeast cells were grown to an optical density (OD) of 12-16 to accumulate inactive ribosome. The cells were harvested and lysed by a standard cryogenic grinding method. Broken cells were thawed on ice and resuspended in a lysis buffer and cleared by centrifugation at 10000 g for 10 min at 4 °C. The resulting supernatant was loaded on top of 10%-50% sucrose gradients and then centrifuged at 28000 rpm in a sw28 rotor for 2.5 hours at 4 °C. Fifty fractions at 800 μl volume were taken from the top to bottom using pipette. The UV (254 nm) absorbance of each fraction was measured using Nanodrop2000. The fractions corresponding to the 80S peak fractions were pooled and pelleted for 1 hour at 436000 g and 4 °C in a TLA-100 rotor. Pelleted 80S were resuspended in the grid preparation buffer and kept on ice until cryo-EM grid preparation. Preliminary reconstruction indicated excellent *nop1*-D243A ribosome but heterogenous *cbf5*-D95A particles purified by this method.

We also purified ribosome by an affinity method in a magnesium-containing buffer that favors binding of the inactive 80S to an IgG column (Rai et al., 2021; Scaiola et al., 2018). The cells were grown similarly as described above to accumulate inactive ribosome and lysed. The resuspended supernatant was incubated with IgG Sepharose beads (GE healthcare) for binding and followed by incubation with a TEV protease treatment mixture. The ribosome eluted from the IgG beads is in good condition for both strains and was used in cryoEM grid preparation in 25mM HEPES-Na pH 7.5, 100 mM NaCl, 10 mM MgCl_2_ and 1 mM dithiothreitol (DTT) and reconstruction.

### Cryo-EM Sample Preparation, Cryo-EM Data Collection and Analysis

5 μl of each ribosome sample at 2-4 units of UV 260nm absorbance was applied to a plasma cleaned 300 mesh UltraAufoil 1.2/1.3 grid (Quantifoil, Germany). The grids were blotted for 2-3 sec at 8 °C in 100% humidity (blot force 1) and plunged into liquefied ethane cooled with liquid nitrogen using Vitrobot MK IV (FEI, Hillsboro OR). The grids were transferred into a Titan Krios (Thermo Fisher) electron microscope equipped with a K3 detector (Gatan) that operated at 300 kV. Micrographs were collected in a movie-mode (frames) using Leginon (Suloway et al., 2005) with a total dose of 60 e^-^/Å^2^ spreading over 70 to 74 frames at a nominal magnification of 81,000X in counting mode to a calibrated pixel size of 1.074 Å/pixel and with a defocus ranging of -1.3 to -2.5 microns. Micrograph was inspected manually and the micrographs showing poor ice, too much drift, bad sample quality, empty hole or signs of astigmatism were discarded. The FSC was estimated at 0.143-cutoff and the local resolution was estimated using ResMap (Kucukelbir et al., 2014).

For the *nop1*-D243A ribosome, a total of 6,460 micrographs were collected, among which, 4,962 micrographs were selected for further processing (Supplementary Figure 2). Frame alignment and dose compensation were carried out with Motincor2 (Zheng et al., 2017) and the Contrast Transfer Function (CTF) parameters were estimated with Gctf (Zhang, 2016) using a wrapper provided in Relion 3.1 (Zivanov et al., 2018). In total, 695,940 particles were picked using the LoG-based autopicking (bin by 4) and divided into two halves (432,895 and 264,100) to perform 2D class averaging and initial 3D classifications. 459,726 particles were further selected based on 2D classes by eliminating bad classes. The initial orientations were obtained via 3D classifications using an 80 Å low-pass filtered 80S map. Particles were further eliminated from the classes resulted in poorly defined maps to a final set of 326,455 particles. The resulting particles were re-extracted without binning and refined to an overall resolution of 3.24 Å. After performing the multiple rounds of CTF refinement as implemented in Relion 3.1, the resolution could be improved to 2.95Å. Additional classification without alignment was performed to further eliminate particles to 297,800 that were re-refined followed by particle polishing to an overall resolution to 2.84 Å. The 2.84 Å-resolution map showed excellent quality for 60S but diffused features for 40S. Focused refinement was subsequently applied to obtain best features for both two subunits. To obtain 60S density, 3D classification without alignment was performed with a customized mask around 60S. The classes showing good 60S density were pooled and re-refined using the 60S mask to the final overall resolution of 2.57 Å (0.143 FSC). To obtain 40S density, 3D classification without alignment was performed by applying customize mask around 40S. The classes showing good 40S density were (289,345 particles) pooled and re-refined using the 40S mask to a final overall resolution of 2.76 Å. We further applied the similar focused refinement procedure on the head and body of the small subunit, respectively leading to final overall resolutions of 2.72 Å for both head and body (0.143 FSC). To sort out Stm1-containing ribosome, particles with good 40S density were classified without alignment using a spherical mask around the mRNA entry channel, leading to 209,566 particles that resulted in good Stm1 density. The Stm1-containing particle set was used in ribosome conformational analysis by 3D classification and multi-body analysis as implemented in Relion 3.1 (Supplementary Figure 2).

For the Cbf5-D95A sample, a total of 6,358 micrographs were collected, among which 5,800 micrographs were selected for further processing. Frame alignment, dose compensation, CTF-estimation and particle picking were carried out as described above (Supplementary Figure 3). The total 726,857 particles were prepared into a stack that was exported to cryoSPARC (Punjani et al., 2017). Multiple rounds of 2D classification and selection resulted in 462,537 good particles. Heterogenous refinement led to five classes among which three were pooled (404,065 particles) and refined in RELION 3.1 leading to a map with well-resolved 60S but diffused 40S density features with an overall resolution of 3.17 Å. The per-particle CTF refinement followed by focus refinement using the 3D custom masks around 60S region and 40S, respectively leading to 2.89 Å and 3.03 Å for the 60S and 40S, respectively. Additional focused refinement on the head and body of 40S subunit using 3D customized masks yielded the resolution of 3.03 Å and 2.99 Å for the head and body, respectively. To clearly identify Stm1-containing ribosome, the particles with the best 40S subunit resolution were classified without alignment into five classes. The predominant class (239,637 particles) was further refined using a custom mask on a 40S subunit resulted in a resolution of 3.36 Å. 3D classification without alignment using a spherical mask around the mRNA entry channel identified two major classes that showed clear Stm1 density and were pooled together for subsequent conformational analysis (Supplementary Figure 3).

### Model building

The X-ray crystal structure of Stm1-bound 80S (PDB ID: 4V88) was used as the initial structure for model building. The 4V88 structure was fit into a composite cryo-EM density map comprising those for the 60S, head and body of 40S subunits. Manual examination and adjustment of the entire structure was carried out before initial rounds of real-space refinement as implemented in PHENIX (Liebschner et al., 2019). Metal ions were checked against the density and the potential coordinate ligands. Sites of methylation were identified by the presence of protruding density continuous from the rest of the chemical moiety and those of pseudouridylation by the presence of hydrogen bond acceptor (mostly water) near the N1 atom of pseudouridine. The presence of modifications is checked against the snoRNA (https://people.biochem.umass.edu/fournierlab/snornadb/mastertable.php) database (Piekna-Przybylska et al., 2007) and the literature (Sloan et al., 2017). Modified nucleotides and protein residues were build using PYMOL (DeLano) while the restraint parameters were generated by JLIGAND (Lebedev et al., 2012) for PHENIX refinement and COOT (Emsley et al., 2010) display. The final models were iteratively refined with the real-space refinement module in PHENIX and manual checking/remodeling in COOT until the best map correlation coefficients and geometric values were reached (Table S2).

### Structure Comparisons

We compared any given pair of densities with UCSF Chimera (Pettersen et al., 2004) in following manner. The two 80S densities segmented by the watershed segmentation feature available in UCSF Chimera (Pettersen et al., 2004) were both aligned against a reference 60S density. The bodies of the 40S in both 80S densities were compared by displaying the location and degree of rotation of the transformation (ratchet) with the command “measure rotation”. Subsequently, the bodies of the 40S in both 80S densities were aligned against a reference body density. The segmented heads of the 40S in the 80S densities were compared by displaying the location and degree of rotation of the transformation (swivel) with the same command.

### Multi-body Analysis

Multi-body refinement was carried out using Relion-3.1 (Nakane et al., 2018) by defining three bodies (60S, 40S body, and 40S head) with a rotation range of 10 degrees in the order of 40S body to 60S and then 40S head to 40S body. Top three represented eigenvectors were analyzed for a given structure and each is are shown as morphed movies (Supplementary Movies 1-6) as well as a set of transformation axes described above between the two extreme maps within each eigenvector (Supplementary Figure 6).

## Deposition of Electron Cryomicroscope Data

The atomic coordinates and associated density maps have deposited at Protein Data Bank with accession codes 7MPJ & EMD-23935 for the *nop1*-D243 ribosome and 7MPI & EMD-23934 for the *cbf5*-D95A ribosome, respectively.

## Acknowledgments

This work was supported by NIH grant R01 GM124622 to H.L. The authors also acknowledge the use of instruments at the Biological Science Imaging Resource supported by Florida State University. The Titan was funded from NIH grant S10 RR025080. The BioQuantum/K3 was funded from NIH grant U24 GM116788. The Vitrobot Mk IV was funded from NIH grant S10 RR024564. The Solaris Plasma Cleaner was funded from NIH grant S10 RR024564. The DE-64 was funded from from NIH grant U24 GM116788.

## Author Contributions

Y.Z. and H.L. designed all experiments. Y.Z. constructed all yeast strains, performed polysome profiling, and purified ribosomes. Y.Z. and J.R. prepared cryoEM grids and collected data. J.R. performed cryoEM analysis with the assistance of Y.Z.. Y.Z., J.R. and H.L analyzed data, wrote and edited manuscript and made figures.

## List of Supplementary materials

**Supplementary Table 1.**
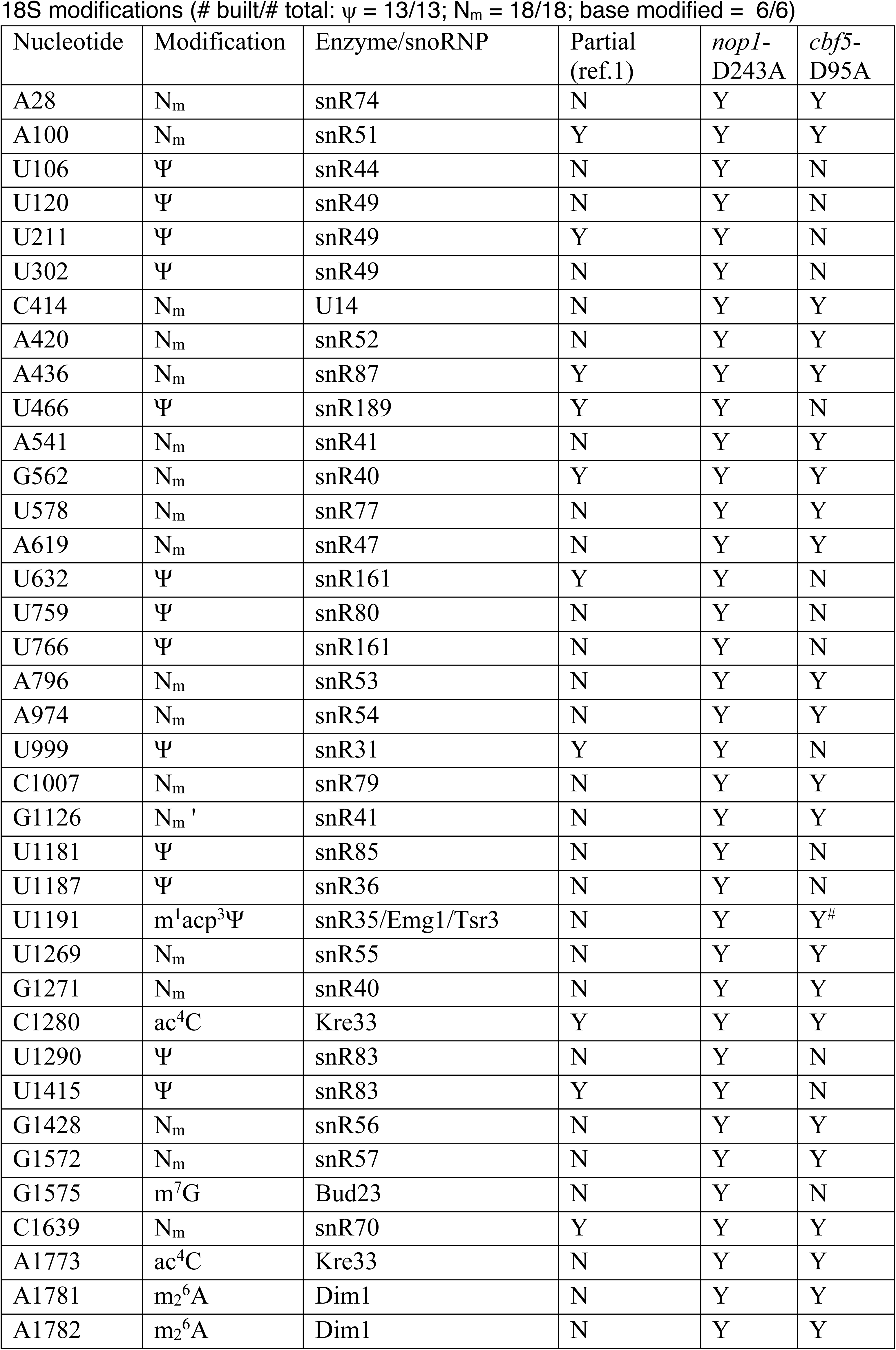
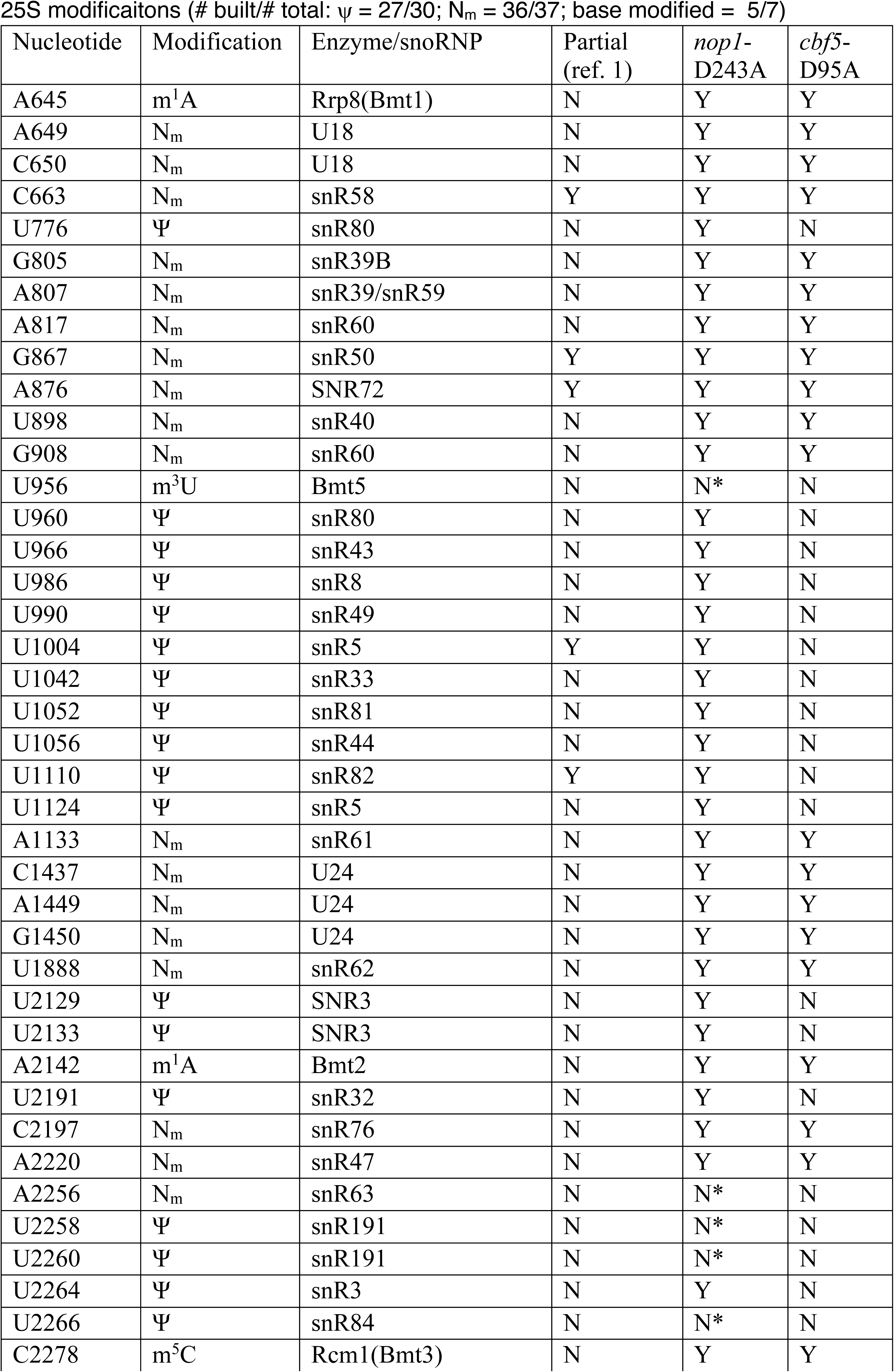
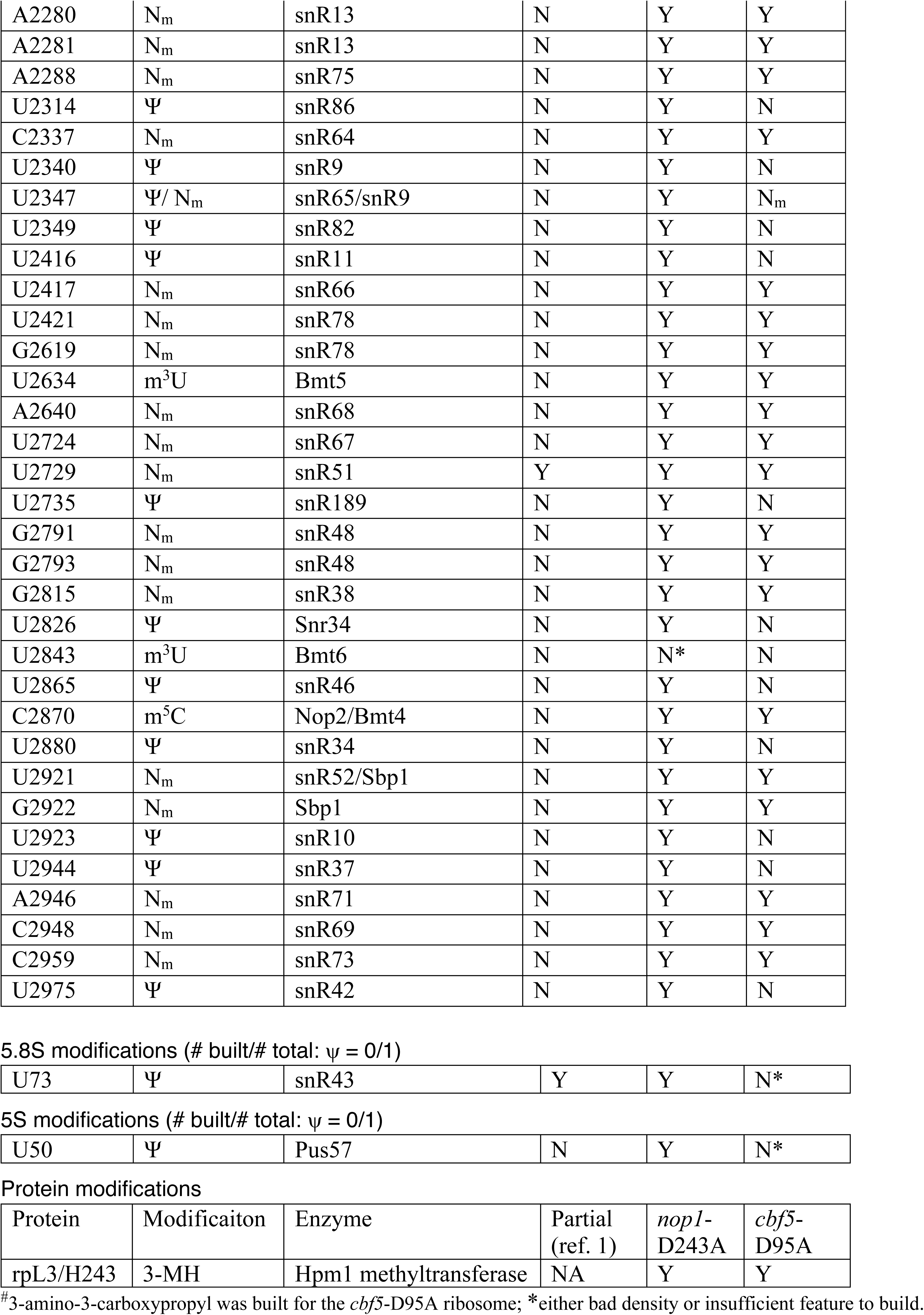
List of Modifications for the *nop1*-D243A and *cbf5*-D95A Ribosome.

modifications (# built/# total: $\psi$ = 13/13; N<sub>m</sub> = 18/18; base modified = 6/6)
| Nucleotide | Modification | Enzyme/snoRNP | Partial (ref.1) | *nop1*-D243A | *cbf5*-D95A |
| --- | --- | --- | --- | --- | --- |
| A28 | N<sub>m</sub> | snR74 | N | Y | Y |
| A100 | N<sub>m</sub> | snR51 | Y | Y | Y |
| U106 | Ψ | snR44 | N | Y | N |
| U120 | Ψ | snR49 | N | Y | N |
| U211 | Ψ | snR49 | Y | Y | N |
| U302 | Ψ | snR49 | N | Y | N |
| C414 | N<sub>m</sub> | U14 | N | Y | Y |
| A420 | N<sub>m</sub> | snR52 | N | Y | Y |
| A436 | N<sub>m</sub> | snR87 | Y | Y | Y |
| U466 | Ψ | snR189 | Y | Y | N |
| A541 | N<sub>m</sub> | snR41 | N | Y | Y |
| G562 | N<sub>m</sub> | snR40 | Y | Y | Y |
| U578 | N<sub>m</sub> | snR77 | N | Y | Y |
| A619 | N<sub>m</sub> | snR47 | N | Y | Y |
| U632 | Ψ | snR161 | Y | Y | N |
| U759 | Ψ | snR80 | N | Y | N |
| U766 | Ψ | snR161 | N | Y | N |
| A796 | N<sub>m</sub> | snR53 | N | Y | Y |
| A974 | N<sub>m</sub> | snR54 | N | Y | Y |
| U999 | Ψ | snR31 | Y | Y | N |
| C1007 | N<sub>m</sub> | snR79 | N | Y | Y |
| G1126 | N<sub>m</sub> ' | snR41 | N | Y | Y |
| U1181 | Ψ | snR85 | N | Y | N |
| U1187 | Ψ | snR36 | N | Y | N |
| U1191 | m<sup>1</sup>acp<sup>3</sup>Ψ | snR35/Emg1/Tsr3 | N | Y | Y<sup>#</sup> |
| U1269 | N<sub>m</sub> | snR55 | N | Y | Y |
| G1271 | N<sub>m</sub> | snR40 | N | Y | Y |
| C1280 | ac<sup>4</sup>C | Kre33 | Y | Y | Y |
| U1290 | Ψ | snR83 | N | Y | N |
| U1415 | Ψ | snR83 | Y | Y | N |
| G1428 | N<sub>m</sub> | snR56 | N | Y | Y |
| G1572 | N<sub>m</sub> | snR57 | N | Y | Y |
| G1575 | m<sup>7</sup>G | Bud23 | N | Y | N |
| C1639 | N<sub>m</sub> | snR70 | Y | Y | Y |
| A1773 | ac<sup>4</sup>C | Kre33 | N | Y | Y |
| A1781 | m<sub>2</sub><sup>6</sup>A | Dim1 | N | Y | Y |
| A1782 | m<sub>2</sub><sup>6</sup>A | Dim1 | N | Y | Y |

25S modificaitons (# built/# total: $\psi$ = 27/30; $N_m$ = 36/37; base modified = 5/7)
| Nucleotide | Modification | Enzyme/snoRNP | Partial<br>(ref. 1) | *nop1*-<br>D243A | *cbf5*-<br>D95A |
| --- | --- | --- | --- | --- | --- |
| A645 | m<sup>1</sup>A | Rrp8(Bmt1) | N | Y | Y |
| A649 | N<sub>m</sub> | U18 | N | Y | Y |
| C650 | N<sub>m</sub> | U18 | N | Y | Y |
| C663 | N<sub>m</sub> | snR58 | Y | Y | Y |
| U776 | Ψ | snR80 | N | Y | N |
| G805 | N<sub>m</sub> | snR39B | N | Y | Y |
| A807 | N<sub>m</sub> | snR39/snR59 | N | Y | Y |
| A817 | N<sub>m</sub> | snR60 | N | Y | Y |
| G867 | N<sub>m</sub> | snR50 | Y | Y | Y |
| A876 | N<sub>m</sub> | SNR72 | Y | Y | Y |
| U898 | N<sub>m</sub> | snR40 | N | Y | Y |
| G908 | N<sub>m</sub> | snR60 | N | Y | Y |
| U956 | m<sup>3</sup>U | Bmt5 | N | N* | N |
| U960 | Ψ | snR80 | N | Y | N |
| U966 | Ψ | snR43 | N | Y | N |
| U986 | Ψ | snR8 | N | Y | N |
| U990 | Ψ | snR49 | N | Y | N |
| U1004 | Ψ | snR5 | Y | Y | N |
| U1042 | Ψ | snR33 | N | Y | N |
| U1052 | Ψ | snR81 | N | Y | N |
| U1056 | Ψ | snR44 | N | Y | N |
| U1110 | Ψ | snR82 | Y | Y | N |
| U1124 | Ψ | snR5 | N | Y | N |
| A1133 | N<sub>m</sub> | snR61 | N | Y | Y |
| C1437 | N<sub>m</sub> | U24 | N | Y | Y |
| A1449 | N<sub>m</sub> | U24 | N | Y | Y |
| G1450 | N<sub>m</sub> | U24 | N | Y | Y |
| U1888 | N<sub>m</sub> | snR62 | N | Y | Y |
| U2129 | Ψ | SNR3 | N | Y | N |
| U2133 | Ψ | SNR3 | N | Y | N |
| A2142 | m<sup>1</sup>A | Bmt2 | N | Y | Y |
| U2191 | Ψ | snR32 | N | Y | N |
| C2197 | N<sub>m</sub> | snR76 | N | Y | Y |
| A2220 | N<sub>m</sub> | snR47 | N | Y | Y |
| A2256 | N<sub>m</sub> | snR63 | N | N* | N |
| U2258 | Ψ | snR191 | N | N* | N |
| U2260 | Ψ | snR191 | N | N* | N |
| U2264 | Ψ | snR3 | N | Y | N |
| U2266 | Ψ | snR84 | N | N* | N |
| C2278 | m<sup>5</sup>C | Rcm1(Bmt3) | N | Y | Y |
| A2280 | N<sub>m</sub> | snR13 | N | Y | Y |
| A2281 | N<sub>m</sub> | snR13 | N | Y | Y |
| A2288 | N<sub>m</sub> | snR75 | N | Y | Y |
| U2314 | Ψ | snR86 | N | Y | N |
| C2337 | N<sub>m</sub> | snR64 | N | Y | Y |
| U2340 | Ψ | snR9 | N | Y | N |
| U2347 | Ψ/ N<sub>m</sub> | snR65/snR9 | N | Y | N<sub>m</sub> |
| U2349 | Ψ | snR82 | N | Y | N |
| U2416 | Ψ | snR11 | N | Y | N |
| U2417 | N<sub>m</sub> | snR66 | N | Y | Y |
| U2421 | N<sub>m</sub> | snR78 | N | Y | Y |
| G2619 | N<sub>m</sub> | snR78 | N | Y | Y |
| U2634 | m<sup>3</sup>U | Bmt5 | N | Y | Y |
| A2640 | N<sub>m</sub> | snR68 | N | Y | Y |
| U2724 | N<sub>m</sub> | snR67 | N | Y | Y |
| U2729 | N<sub>m</sub> | snR51 | Y | Y | Y |
| U2735 | Ψ | snR189 | N | Y | N |
| G2791 | N<sub>m</sub> | snR48 | N | Y | Y |
| G2793 | N<sub>m</sub> | snR48 | N | Y | Y |
| G2815 | N<sub>m</sub> | snR38 | N | Y | Y |
| U2826 | Ψ | Snr34 | N | Y | N |
| U2843 | m<sup>3</sup>U | Bmt6 | N | N* | N |
| U2865 | Ψ | snR46 | N | Y | N |
| C2870 | m<sup>5</sup>C | Nop2/Bmt4 | N | Y | Y |
| U2880 | Ψ | snR34 | N | Y | N |
| U2921 | N<sub>m</sub> | snR52/Sbp1 | N | Y | Y |
| G2922 | N<sub>m</sub> | Sbp1 | N | Y | Y |
| U2923 | Ψ | snR10 | N | Y | N |
| U2944 | Ψ | snR37 | N | Y | N |
| A2946 | N<sub>m</sub> | snR71 | N | Y | Y |
| C2948 | N<sub>m</sub> | snR69 | N | Y | Y |
| C2959 | N<sub>m</sub> | snR73 | N | Y | Y |
| U2975 | Ψ | snR42 | N | Y | N |
| 5.8S modifications (# built/# total: ψ = 0/1) |  |  |  |  |  |
| U73 | Ψ | snR43 | Y | Y | N* |
| 5S modifications (# built/# total: ψ = 0/1) |  |  |  |  |  |
| U50 | Ψ | Pus57 | N | Y | N* |

Protein modifications
| Protein | Modificaiton | Enzyme | Partial<br>(ref. 1) | *nop1*-<br>D243A | *cbf5*-<br>D95A |
| --- | --- | --- | --- | --- | --- |
| rpL3/H243 | 3-MH | Hpm1 methyltransferase | NA | Y | Y |

<sup>#</sup>3-amino-3-carboxypropyl was built for the *cbf5*-D95A ribosome; \*either bad density or insufficient feature to build.

**Supplementary Table 2.** Data Acquisition, Processing and Model Refinement.

| **Data acquisition & processing parameters** | ***nop1*-D243A** | ***cbf5*-D95A** |
| --- | --- | --- |
| Microscope | Titan Krios | Titan Krios |
| Detector | Gatan K3 | Gatan K3 |
| Voltage | 300 kV | 300 kV |
| Electron Source | Field Emission Gun | Field Emission Gun |
| Collecting Mode | Counting | Counting |
| Dose Rate (e<sup>-</sup>/Å<sup>2</sup>) | 60.07 | 60.07 |
| Defocus range (μm) | -1.3 to -2.5 | -1.3 to -2.5 |
| Nominal Magnification | 81000X | 81000X |
| Frames collected per exposure | 70 | 74 |
| Framealignment Software | MotionCor2 | MotionCor2 |
| CTF parameter estimation | Gctf | Gctf |
| Total number of raw images collected | 6,460 | 6,358 |
| Number of images used for particle picking | 4,962 | 5,800 |
| Initial particles picked | 695,940 | 726,857 |
| 2D classification software | Relion 3.1 | Cryosparc |
| Final reconstruction software | Relion 3.1 | Relion 3.1 |
| Applied symmetry | C1 | C1 |
| Number of particles contributed for the final reconstruction | 288,240 (60S)<br>289,345 (40S) | 404,065 (60S)<br>349,046 (40S) |
| Resolution method | FSC 0.143 cut-off | FSC 0.143 cut-off |
| Map resolution (Å) | 2.57 Å (60S)<br>2.72 Å (40S-head)<br>2.72 Å (40S-body) | 2.89 Å (60S)<br>3.03 Å (40S-head)<br>2.99 Å (40S-body) |
| Local resolution determining Software | Resmap | Resmap |
| Map Visualization software | Pymol/Chimera/<br>Chimera X/ Coot | Pymol/Chimera/<br>Chimera X/ Coot |
| Deposit EMDB codes | 23935 | 23934 |
| **Refinement parameters** |  |  |
| CC (map_model) | 0.78 (mask), 0.81 (volume) | 0.78 (mask), 0.82 (volume) |
| RMSD (Bond lengths/Bond angles) | 0.012/1.222 | 0.009/1.096 |
| Ramachandran plot (%) (Outlier/Allowed/Favored) | 0.45/7.81/91.74 | 0.3/8.25/91.35 |
| Cβ Outliers (%) | 0.03 | 0.02 |
| MolProbity Score | 1.71 | 1.81 |
| Clash score | 4.29 | 5.52 |
| Rotamer outliers (%) | 0.05 | 0.03 |
| ADP (B-factor) | 199811 | 199433 |
| Protein (min/max/mean) | 2.57/32.30/15.34 | 9.42/104.99/40.21 |
| Nucleotide (min/max/mean) | 4.62/61.43/19.14 | 6.07/183.35/49.97 |
| Ligand (min/max/mean) | 5.63/40.73/12.89 | 7.44/110.30/31.20 |
| dFSC model (0.5) | 2.7 | 2.9 |
| Deposit PDB codes | 7MPJ | 7MPI |

**Supplementary Figure 1.**
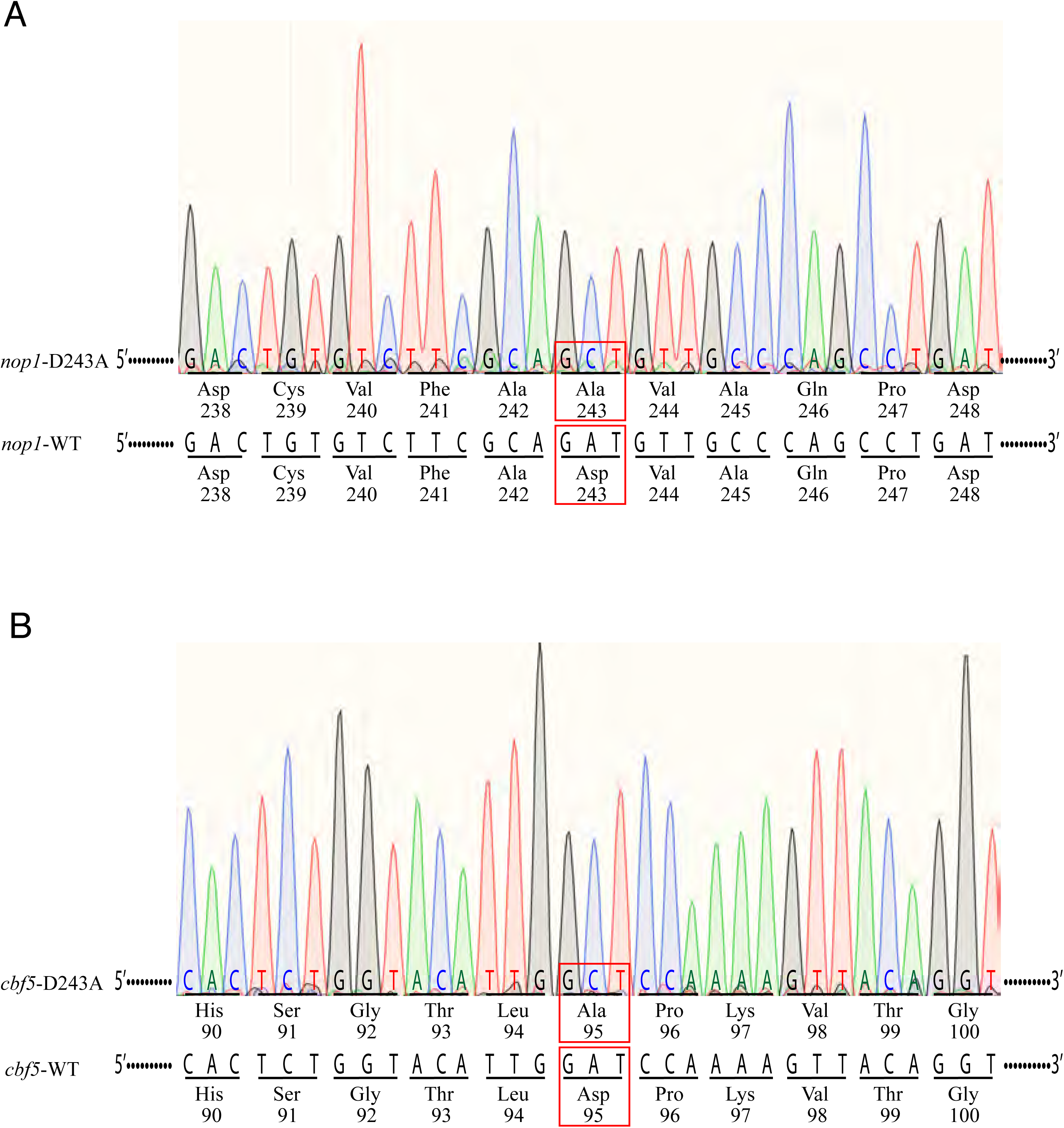
Related to Figures 1, 2, 3, & 4. Sanger sequencing verification of the *cbf5*-D95A (A) and *nop1*-D243A (B) mutations. Interpreted bases from chromatograms are shown in colored letters. Corresponding codons are underscored and the encoded amino acids are show as three-letter symbols. The wild-type sequences of the same regions are shown in black. Point mutations are highlighted with red boxes.

**Supplementary Figure 2.**
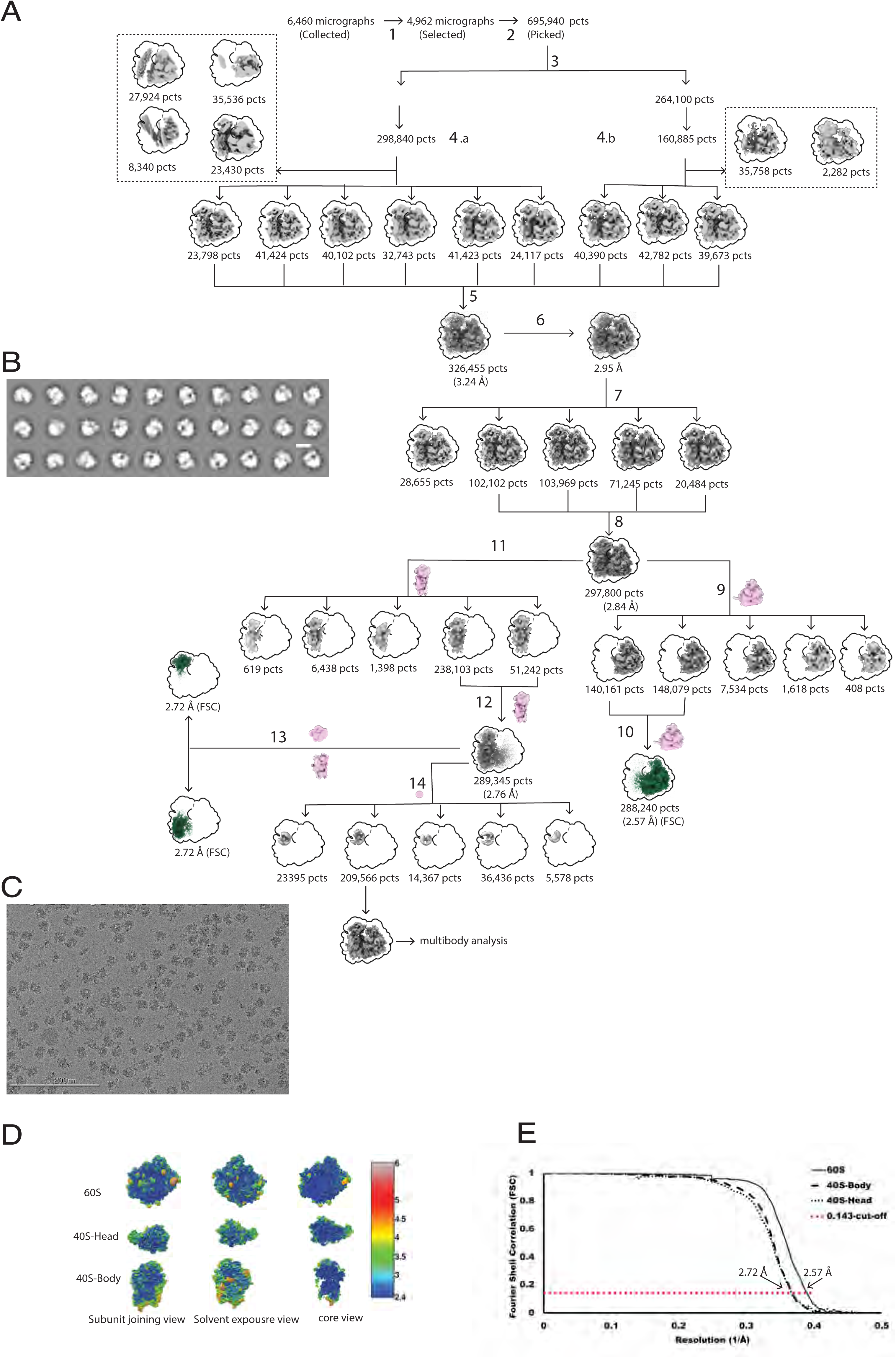
Related to Figures 2, 3, & 4. Cryo-EM image analysis flow chart of *nop1*-D243A ribosome. **A.** Image selection, particle selection, and recontruction schemes. Masks used in focused classification and refinement are shown in pink and at each relevant path junction. The final maps used for model building and refinement are shown in blue. A final of 4,962 images were selected for further processing (1) and a total of 695,940 particles (pcts) were picked (3), extracted in bin 4, and split arbitrarily into two halves. The first half contained 431,895 pcts which were reduced to 298,840 pcts after 2D classification (4.a). The other half contained 264,100 pcts which was reduced to 160,885 pcts after 2D classification (4.b). Particles with bad features shown in dash line boxes were removed. 3D classification of the two half particles resulted in a total of 326,455 pcts that were reextracted in bin 1 (unbinned) and refined to 3.24 Å (5). Contrast Transfer Function (CTF) refinement improved the overall resolution to 2.95 A (6). Further classifification without alignment allowed additional selection of particles (7) for refinment and particle polishing, which led to an overall resolution of 2.84 Å (8). To improve the 60S density, particles were further classified without alignment into five classes using a 3D custom mask on the 60S subunit region (9). The classes with good 60S density were pooled together and again refined using a custom3D mask around the 60S subunits that resulted in an overall resolution of 2.57 Å for the 60S region (10). To improve the 40S density, particles from step 8 were classified without alignment into five classes using the 3D custom mask on the 40S subunit region (11). The classes with good 40S density were pooled and re-refined using a 3D mask around the 40S region that resulted in an overall resolution of 2.76 Å resolution around the 40S region(12). The particles were further refined using a mask around the head and body of the 40S that improved the resolution of head and body to 2.72 Å for both regions (13). To select Stm1-bound ribosome, focused 3D classification was performed using a spherical mask around the mRNA entry channel, which led to a major class with 209,566 particles that showed good Stm1 density and was, therefore, used for multibody analysis (14). **B.** Selected 2D class averages (scale bar 20nm). **C.** A representative raw micrograph (scale bar 200 nm). **D.** Local resolution estimated by Resmap showing the comparably high-resolution inner core. **E.** The Fourier Shell Correlation (FSC) curves of 80S, 40S body, 40S head and 60S. 0.143 cutoff was used for resolution estimatimation.

**Supplementary Figure 3.**
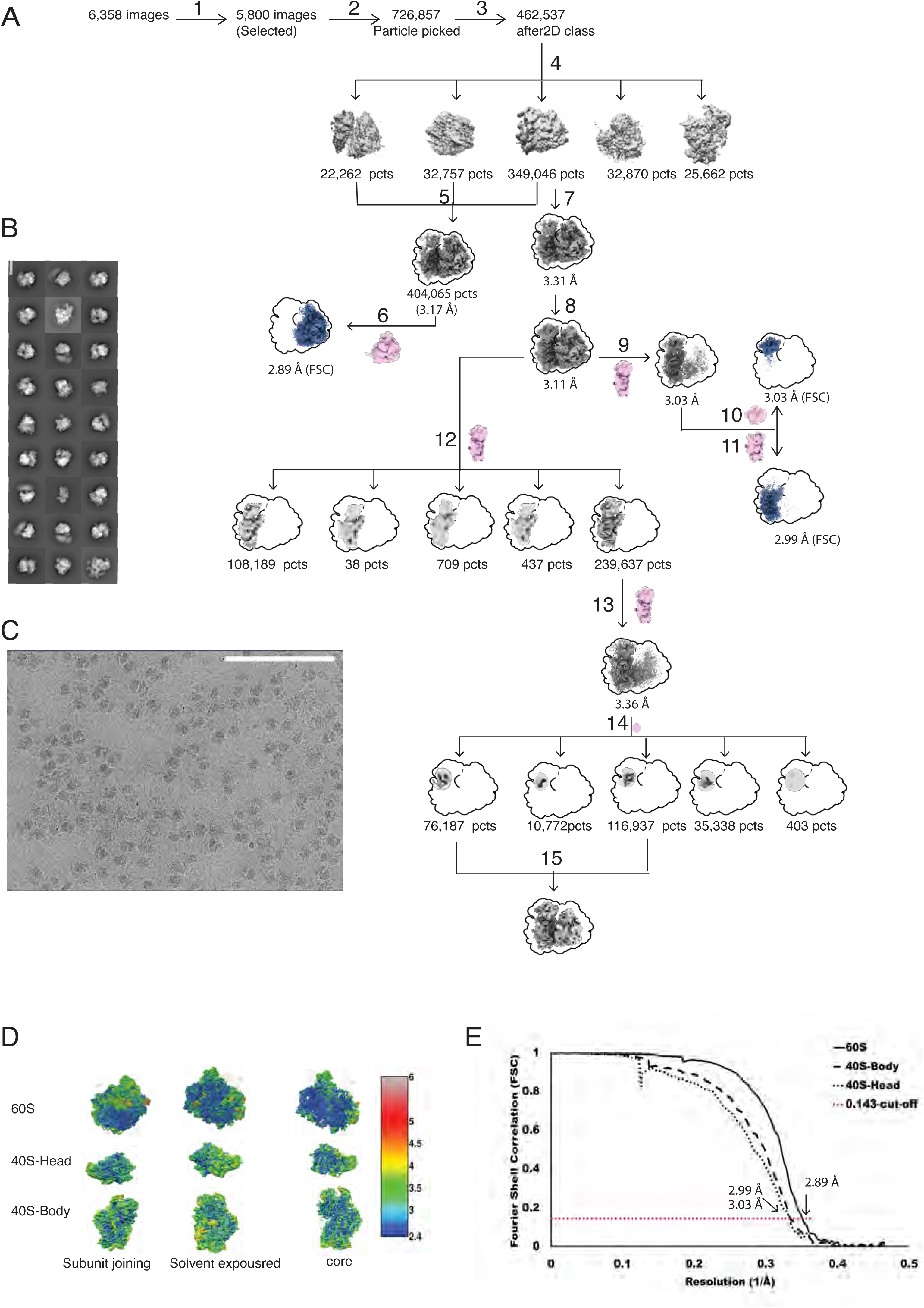
Related to Figures 2, 3, & 4. Cryo-EM image analysis flow chart of *cbf5*-D95A. **A.** Image selection, particle selection, and recontruction schemes. Masks used in focused classification and refinement are shown in pink and at each relevant path junction. The final maps used for model building and refinement are shown in blue. A final of 5800 images were selected for further processing (1) and a total of 726,857 particles (pcts) were picked in RELION 3.1 (2). 2D classifications and selection in cryosparc reduced the particles to 462,537 (3) that were 3D classified into five classes (4). To obtain the best 60S structure, three classes were combined and refined to 3.17 Å in RELION 3.1 (5). After per-particle Contrast Transfer Function (CTF) refinement using the custom mask around the 60S subunit improved the 60S resolution to 2.89 Å (6). To obtain the best 40S resolution, a single dominant class (349,046 pcts) was refined to 3.31 Å resolution (7) which was further improved to 3.11 Å after per particles CTF refinement (8). Using a custom 3D mask on the 40S subunit during refinement yielded the overall 40S subunit resolution to 3.03 Å (9) that was further improved for the body and head region to 3.03 Å (10) and 2.99 Å (11) and 3.03 Å with their respective masks. To ensure multibody analysis was performed on Stm1-bound ribosome, particles with good 40S density were further classified into five classes without alignment using the 40S custom mask (12). The predominant class (239,637 pcts) showed good 40S subunit density and refined using a custom 3D mask on 40S subunit (13) followed by another round of 3D classification based on Stm1 using a spherical mask around the mRNA entry channel (14). Two classes show clear Stm1 density and hence pooled together (15). **B.** Selected 2D class averages (scale bar 20nm). **C.** A representative raw micrograph (scale bar 200 nm). **D.** Local resolution estimated by Resmap showing the comparably high-resolution inner core. **E.** The Fourier Shell Correlation (FSC) curves of 80S, 40S body, 40S head and 60S. 0.143 cutoff was used for resolution estimatimation.

**Supplementary Figure 4.**
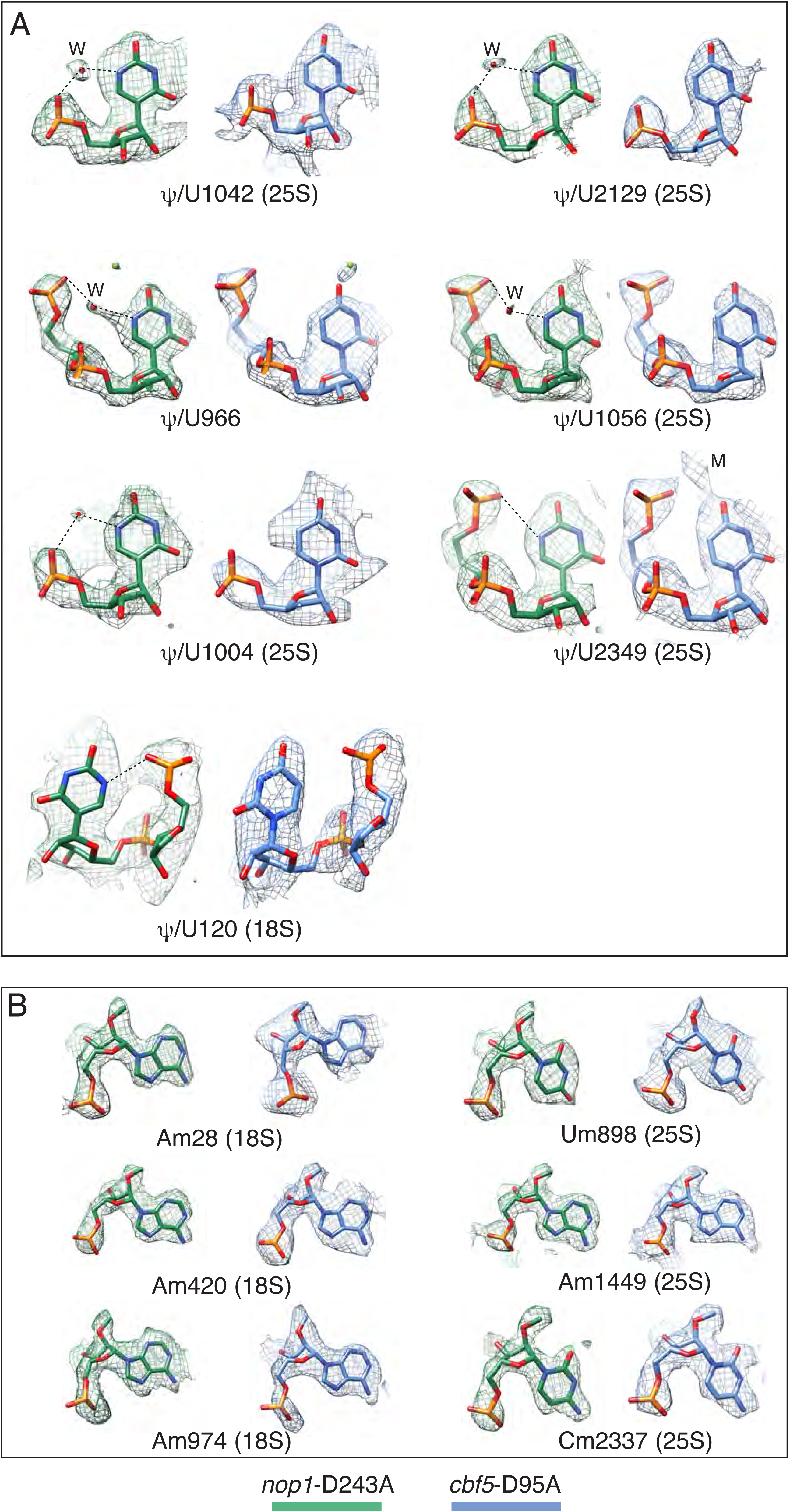
Related to Figures 2, 3, & 4. Electron potential density for selected nucloetides that are predicted to be pseudouridylated (A) or 2’-O-methylated (B). Nucleotides colored in green are those from the *nop1*-D243A ribosome and those colored in blue from from the *cbf5*-D95A ribosome. “W” denotes a potential water molecule and “M” denote a potential metal ion. Dashed lines denote close contacts.

**Supplementary Figure 5.**
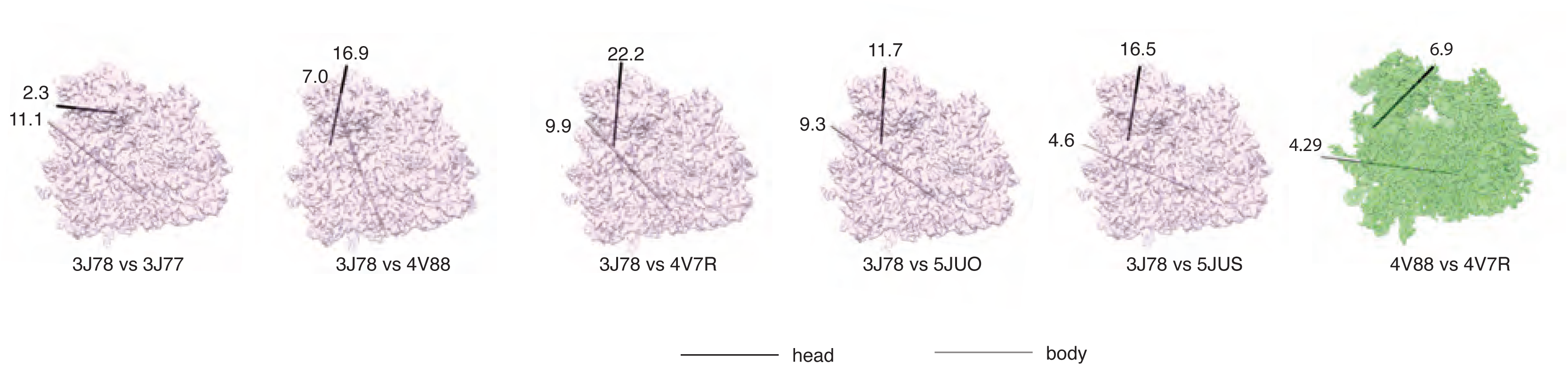
Related to Figure 3. Comparison of known ribosome conformations. For each compared ribosome, Protein Data Bank (PDB) code is given. Numers indicate degrees of rotation in clockwise direction when viewed into each axis. The cryoEM maps (or model converted maps) were used to compute their relative motion with respect to the non-rotated state (PDB ID: 3J78, pink) or between the two Stm1-bound ribosome (green). The relationship between the two compared heads is indicated by the location and degree of rotation of the rotation axis in black (swivel). The relationship between the two compared bodies is indicated by the location and degree of rotation of the rotation axis in gray (ratchet).

**Supplementary Figure 6.**
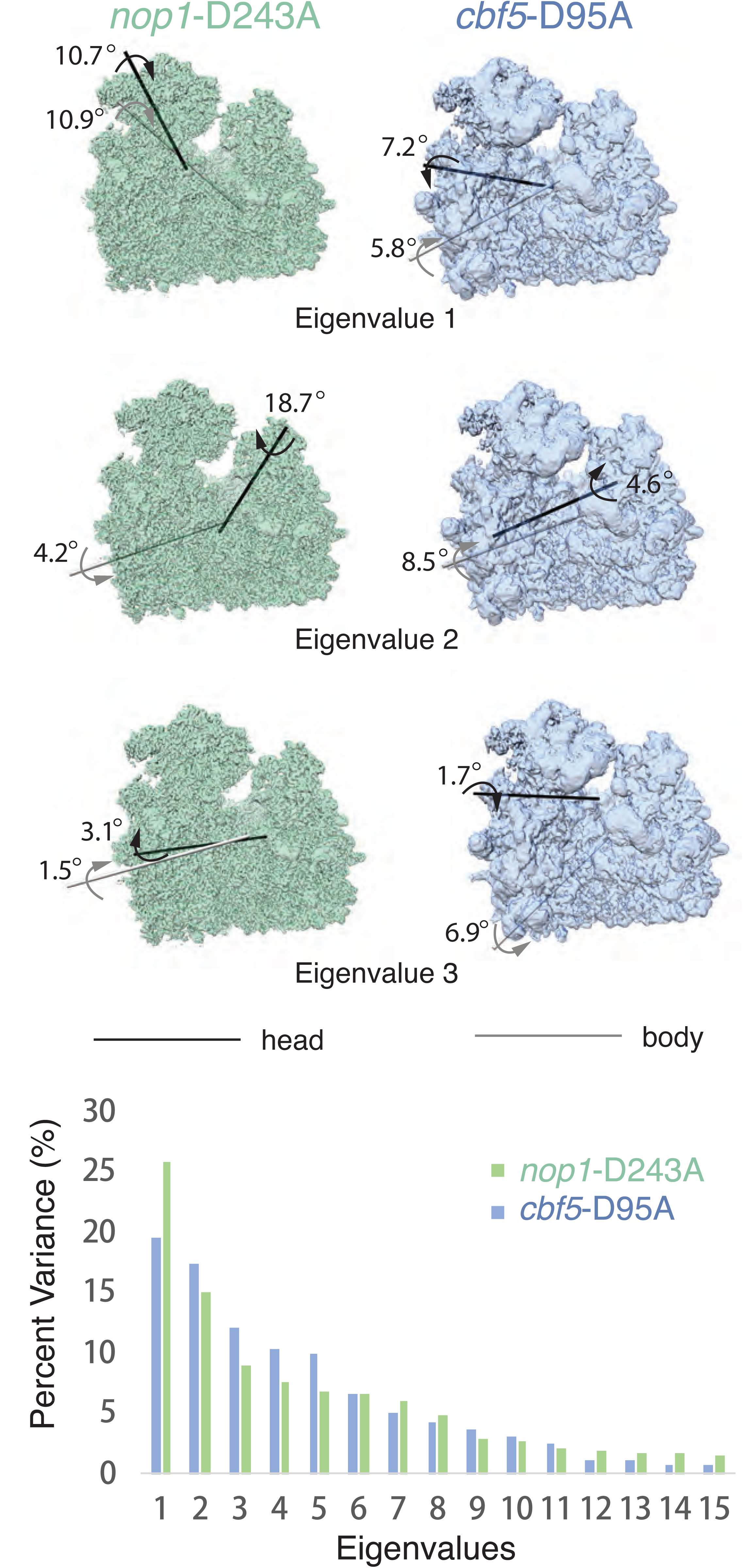
Related to Figure 3. 3D multivaribilty analysis for the *nop1*-D243A (green) and *cbf5*-D95A (blue) ribosome. Top, electron potential densities representing the two extreme amplitude values of top 3 eigenvectors are compared. For each eigenvector, the two extreme maps were superimposed to a reference 60S density in order for the rotation axis that brings the 40S body of one map to that of another map to be calculated. The two maps were then superimposed to a reference map of the 40S body for the rotation axis that brings the 40S head of one map to that of another map to be calculated. The arrows indicate the direction of rotation while the numbers indicate the degree of rotation. Bottom, percent variance plot for the top 15 eigenvectors for the *nop1*-D243A (green) and *cbf5*-D95A (blue) ribosome.

**Supplementary Figure 7.**
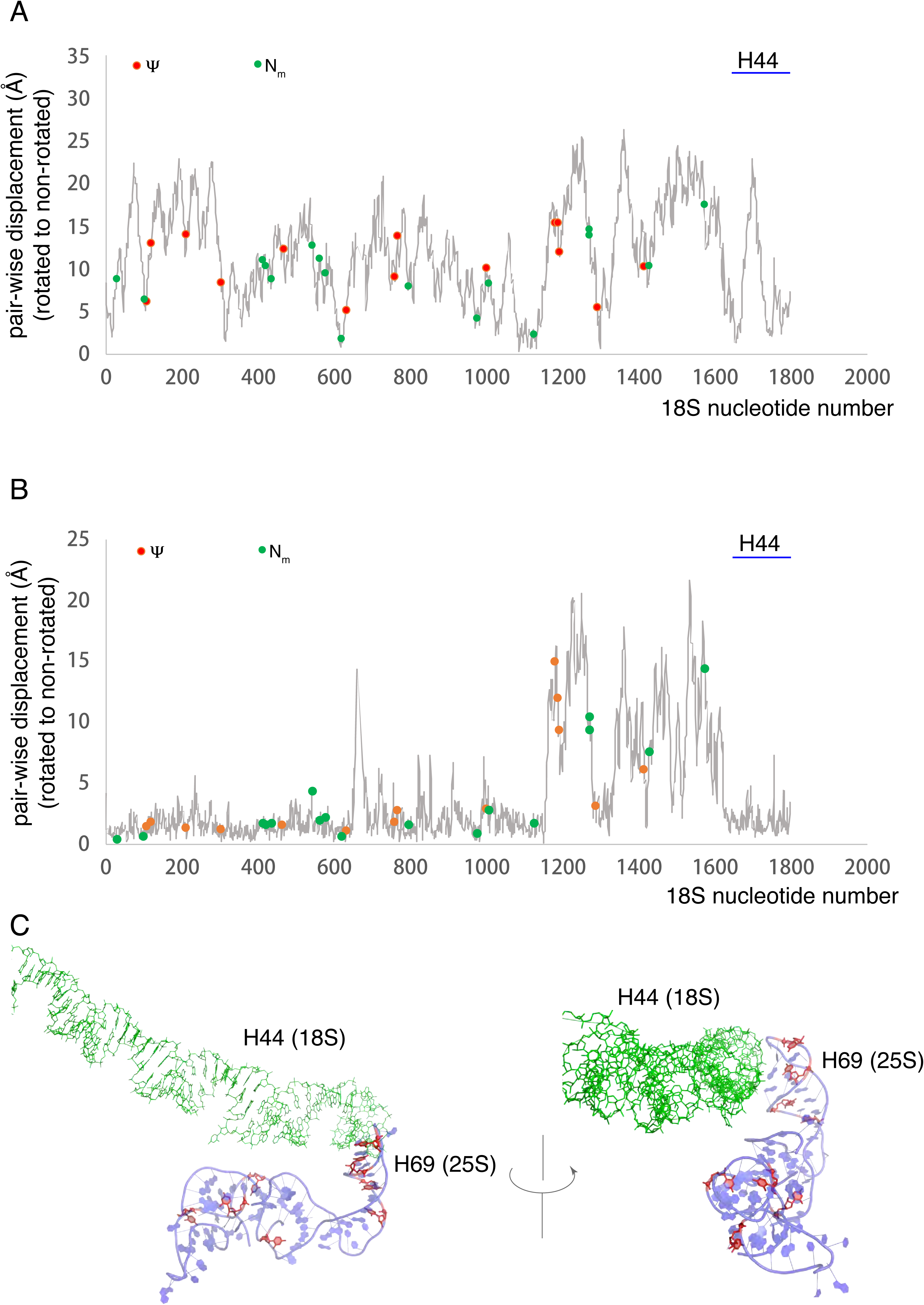
Related to Figure 4. Mapping modification sites on the nucleotide pair-wise displacement plot for the small subunit ribosomal RNA (18S rRNA) when transforming the rotated (PDB ID: 3J77) to the non-rotated (PDB ID: 3J78) ribosome conformation. Red spheres indication positions of pseudouridine while green spheres indicate the positions of 2’-O-methylated nucloetides. The location of helix H44 is indicated by a blue bar. **A**. Pair-wise displacement between 3J77 and 3J78 18S nucleotides when their 25S rRNA are aligned. **B**. Pair-wise displacement between 3J77 and 3J78 18S nucleotides when the body porsion of their 18S rRNA are aligned. **C.** Illustration of interactions between helix H44 of 18S (green) with a pseudouridine-rich helix H69 at the intersubunit bidge region of 25S (blue). H44 nucleotides are shawn as sticks models while H69 is shown as cartoon models. Pseodouridine is colored in red.

**Supplementary Figure 8.**
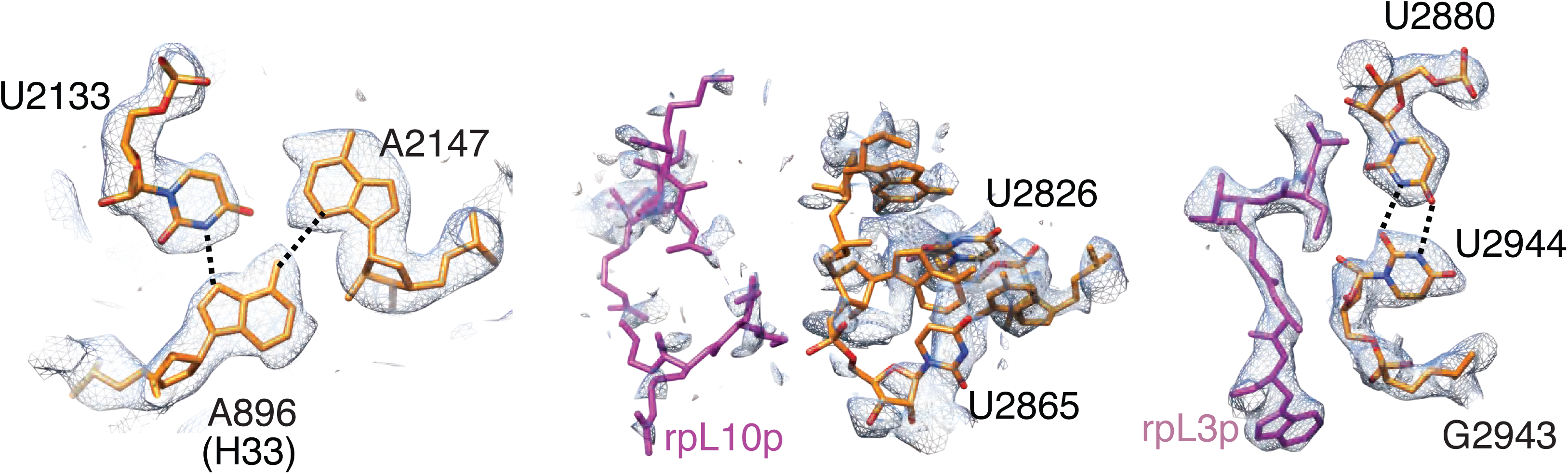
The correspondng electron potential density maps in the *cbf5*-D95A ribosome around the pseodouridine clusters on the large subunit shown in Figure 4C for *nop1*-D243A ribosome.

**Supplementary Movie 1. Related to** Figure 3 **&** Supplementary Figure 6. Domain motions representing eigenvector 1 resulted from multivaribility analysis of *nop1*-D243A ribosome.

**Supplementary Movie 2. Related to** Figure 3 **&** Supplementary Figure 6. Domain motions representing eigenvector 1 resulted from multivaribility analysis of *Cbf5*-D95A ribosome.

**Supplementary Movie 3. Related to** Figure 3 **&** Supplementary Figure 6. Domain motions representing eigenvector 2 resulted from multivaribility analysis of *nop1*-D243A ribosome.

**Supplementary Movie 4. Related to** Figure 3 **&** Supplementary Figure 6. Domain motions representing eigenvector 2 resulted from multivaribility analysis of *Cbf5*-D95A ribosome.

**Supplementary Movie 5. Related to** Figure 3 **&** Supplementary Figure 6. Domain motions representing eigenvector 3 resulted from multivaribility analysis of *nop1*-D243A ribosome.

**Supplementary Movie 6. Related to** Figure 3 **&** Supplementary Figure 6. Domain motions representing eigenvector 3 resulted from multivaribility analysis of *Cbf5*-D95A ribosome.

